# The Fem Cell-surface Signaling System is Regulated by ExsA in *Pseudomonas aeruginosa* and Affects Pathogenicity

**DOI:** 10.1101/2024.06.04.597374

**Authors:** Yanqi Li, Sara Badr, Pansong Zhang, Lin Chen, Anjali Y. Bhagirath, Maryam Dadashi, Michael G. Surette, Kangmin Duan

## Abstract

Bacterial interspecies interactions shape the function and the structural dynamics of microbial communities and affect the progression of polymicrobial infections. The FemA-FemR-FemI (Fem) cell surface signaling system in *Pseudomonas aeruginosa* is known to be involved in the uptake of iron-chelating mycobactin produced by the *Mycobacterium* species. In this report, we present the data that indicates the *femA*-PA1909 operon was positively regulated by ExsA, a master regulator for the type three secretion system (T3SS), connecting the Fem system with T3SS. Intriguingly, the Fem system also influenced virulence factors in *P. aeruginosa,* including the quorum sensing systems, pyocyanin production, biofilm formation and the type six secretion systems (T6SSs). Using a *Galleria mellonella* infection model we observed that an *femA* deletion in PAO1 significantly increased the host survival rate while *femI* over-expression decreased it, suggesting a role for the Fem system in bacterial pathogenicity *in vivo*. Our data indicate the Fem system is a target of the T3SS master activator ExsA, and it affects *P. aeruginosa* pathogenicity.

---

*Pseudomonas aeruginosa* is one of the predominant nosocomial pathogens causing pneumonia, burn wounds, and urinary tract infections in immunocompromised individuals. It is also one of the three key pathogens identified by the World Health Organization (WHO) whose antibiotic resistance poses the highest threat to human health ^1^. In patients with cystic fibrosis (CF), chronic polymicrobial infections caused by *P. aeruginosa* and other bacteria in the respiratory tract and the associated pulmonary inflammation are responsible for the majority of pulmonary failure and eventual mortality ^2^.

Interspecies interactions within the microbial communities associated with polymicrobial diseases play an important role in disease manifestations and outcomes ^3^. A strong association in disease severity has been shown between *P. aeruginosa* and the presence of other lung microbiota in CF ^4^. Complex interactions between *P. aeruginosa* and oropharyngeal microbiota including *Streptococcus* milleri/anginosus groups can significantly affect the disease progression in CF lungs, ^5,6^*. P. aeruginosa, Burkholderia cenocepacia,* and *Stenotrophomonas maltophilia* interact through diffusible signal factor (DSF) signaling, whereas *P. aeruginosa* responds to the DSF produced by *B. cenocepacia,* and *S. maltophilia* by reducing the type three secretion system (T3SS) and modulating biofilm formation respectively ^7^. Nontuberculous mycobacteria (NTM) are emerging pathogens in CF, present in 10% of these patients, and increasingly cause problematic nosocomial infections^8^. However, it is not known if and how *P. aeruginosa* and NTM interact in a microbial community or in CF lungs.

The pathogenic potential of *P. aeruginosa* is attributed to its myriad of virulence factors. One of the major virulence factors in *P. aeruginosa* is T3SS which injects deleterious effector proteins directly from bacterium cytosol into host cells ^9^. The four known T3SS effector proteins (ExoU, ExoY, ExoT, and ExoS) promote host tissue destruction and modulate the inflammatory responses, which enable *P. aeruginosa* to evade phagocytosis and disseminate from initial colonization site ^10^. T3SS expression is strictly controlled by both global and finely-tuned regulatory mechanisms ^11^. The transcription of the T3SS genes is controlled by the master regulator, ExsA, a member of the AraC/XylS family of transcriptional activators, which recognizes and binds to the conserved consensus sequence in the promoters of T3SS genes. Recent studies have indicated that ExsA may regulate genes not part of the T3SS, such as *impA* encoding an extracellular metalloprotease that inhibits phagocytosis of macrophages ^12^. In a genome-wide ChIP-seq analysis of virulence-related transcriptional factors, Huang *et al* revealed that ExsA binds to the promoters of non-T3SS genes, indicating that ExsA is able to regulate bacterial activities other than T3SS ^13^.

Cell-surface signal systems are tripartite molecular devices that allow Gram-negative bacteria to sense extracellular stimuli and transmit them into a coordinated transcriptional response ^14^. Fem system (FemA-FemR-FemI) in *P. aeruginosa* is such a cell-surface signal system that the bacterium uses to acquire iron through mycobactin, a heterogenous siderophore produced by *Mycobacterium* species ^15^. The Fem system consists of an outer-membrane-located TonB-dependent transducer FemA, a cytoplasmic membrane-located anti-sigma factor FemR (PA1911), and an extra-cytoplasmic function (ECF) sigma factor-FemI (PA1912). In the presence of mycobactin, FemA uptakes mycobactin and TonB interacts with the FemA to generate a signal that is transmitted to the anti-sigma factor FemR, causing the release of the sigma factor FemI into the cytoplasm. FemI can bind to the core RNA polymerase (cRNAP) and activate transcription of the *femA* operon ^15^.

In this report, we present data indicating that the PA1910(*femA*)-PA1909 operon in *P. aeruginosa*, referred to as *femA* collectively hereafter, is a target of the T3SS master regulator ExsA. Binding of ExsA to the promoter region of *femA* was confirmed by electrophoretic mobility shift assay (EMSA). Our results also show that the Fem system not only functions as an iron uptake system for heterogenous siderophore, but also affects *P. aeruginosa* activities including *in vivo* pathogenicity, independent of the iron transport function.

## RESULTS

### The Fem system in P. aeruginosa is directly regulated by ExsA, the key regulator of the T3SS

In *P. aeruginosa,* T3SS-related genes are organized in 10 transcriptional units which encode the essential components and regulatory proteins of T3SS ^16^. All these units have the conserved nucleotides in the promoter regions which the T3SS central regulator ExsA recognizes and to which it binds. ExsA monomer first binds to the binding site-1 (**GrCynnnmYTGayAk**) and then recruits a second ExsA to occupy the binding site-2. In addition, occupation of the binding site-2 (**tAaAAA**) is dependent upon monomer-monomer interactions mediated by the amino-terminal domain of ExsA ^11,17^. During analyzing the whole genome sequences of *P. aeruginosa* PAO1, we found that the promoter region of the operon [PA1910 (*femA*)-PA1909], possesses ExsA binding site-1 and binding site-2 (Fig. 1). This led us to investigate the possibility that ExsA binds to the *femA* promoter region and regulates the Fem iron uptake system.

**FIG. 1.**
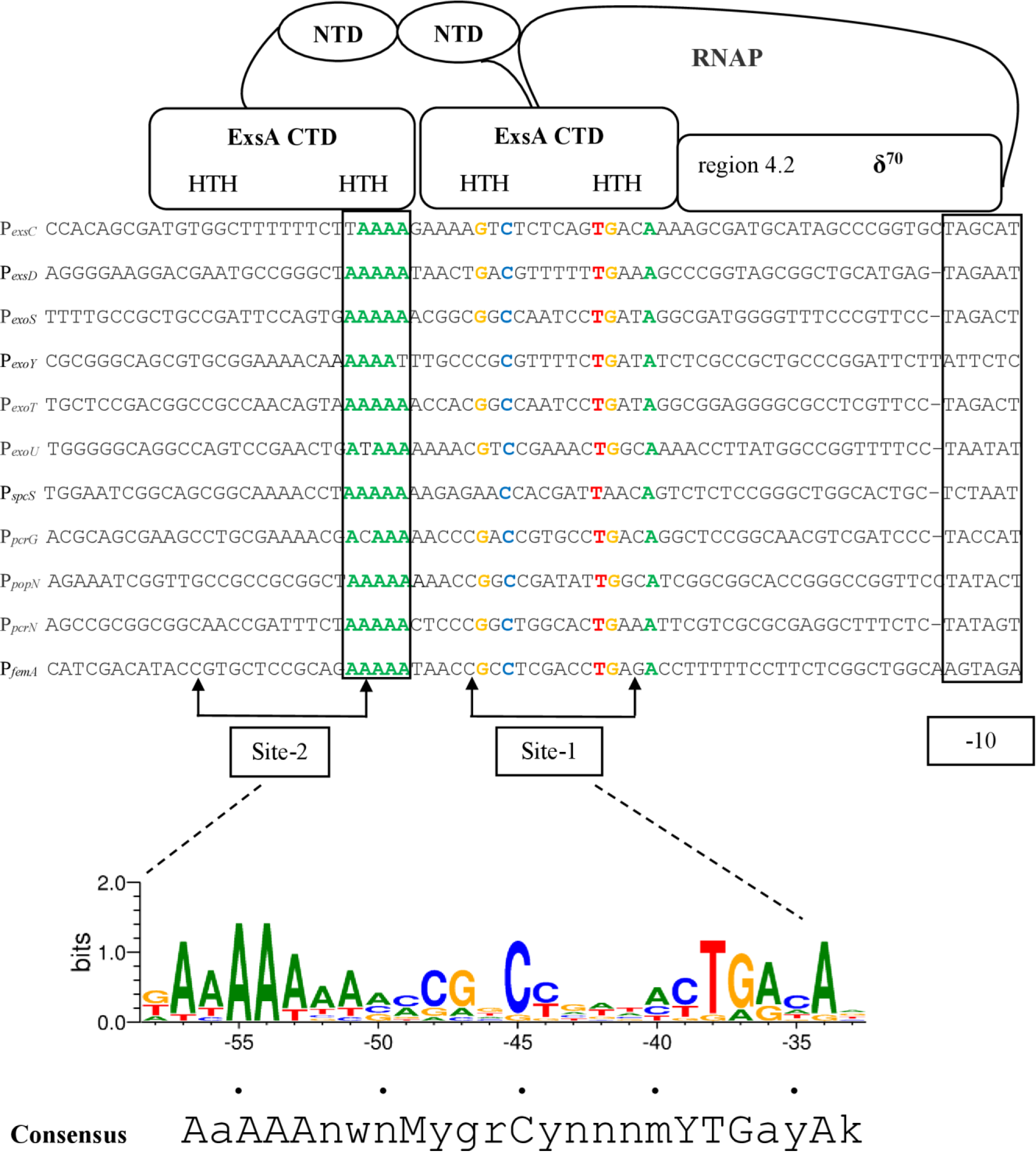
Sequence alignment of ExsA-dependent promoters and the model for promoter recognition and recruitment of σ^70^-RNAP by ExsA. Highly conserved nucleotides in all ExsA-dependent promoters that are essential for ExsA-dependent activation are shown in bold colors. The predicted interaction between the helix-turn-helix (HTH) DNA-binding motifs of each ExsA monomer is shown. ExsA interacts with the region 4.2 of σ^70^ to recruit σ^70^-RNAP. The predicted ExsA consensus binding sequence is indicated at the bottom of the diagram. Figure adapted from Diaz et al. (11).

To examine the interaction between ExsA and promoter of *femA*, electrophoretic mobility shift assay (EMSA) was carried out. ExsA was expressed as an N-terminal histidine-tagged fusion protein (ExsA_His_) in *E. coli* and purified by affinity chromatography. The 6-FAM^TM^-labeled DNA fragment of the *femA* promoter region (∼200 bp) was amplified by PCR. We used the promoter region of *exoT*, an ExsA-regulated gene, as the positive control for the EMSA. Two different shifted bands were observed with the reaction mix of ExsA_His_ and *femA* promoter (Fig. 2). Like previously demonstrated ExsA binding to T3SS promoters ^17^, the higher-mobility ExsA-DNA complex probably represents initial ExsA bound to site 1 in the promoter region and the lower-mobility complex represents ExsA bound at both sites. The binding could be inhibited with an approximately 10-fold concentration of unlabeled *femA* promoter fragment, while no effect was observed upon addition of the same amount of nonspecific unlabeled DNA (*norC*). The results indicate that the ExsA can specifically bind to the *femA* promoter region, presumably regulating the promoter activity.

**FIG. 2.**
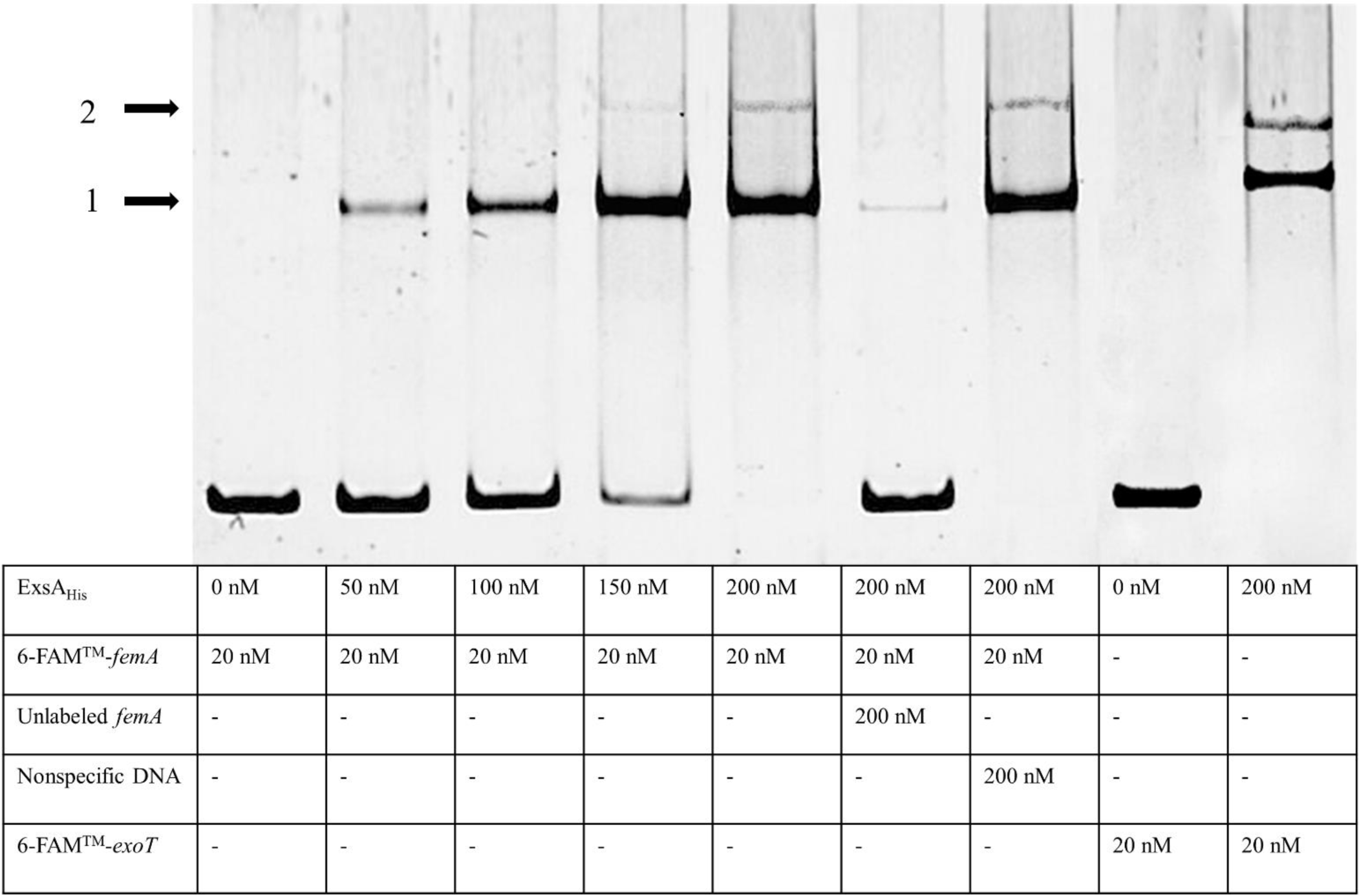
EMSA displaying the binding of ExsA to the *femA* promoter. ExsA was expressed and purified as an amino-terminal histidine-tagged fusion protein (ExsA_His_) in *E. coli* BL21 (DE3). DNA fragment containing the *femA* promoter region (200 bp) was generated by PCR. The protein concentration used in each sample is indicated in the box below. Two shifted bands 1 and 2 are indicated by arrows. Higher-mobility ExsA-DNA complex consisting of a single ExsA_His_ molecule bound to the DNA whereas the lower-mobility complex results from the binding of two ExsA_His_ molecules. DNA fragment containing the *exoT* promoter was used as a positive control; Nonspecific DNA, the promoter region of *norC* that is not an ExsA target was used as unlabelled non-specific competitor DNA.

### The FemA operon is regulated by ExsA, independent of FemI

To verify the regulatory effect of ExsA on the *femA* operon, we compared the promoter activities of the *femA-*PA1909 operon in PAO1 and in an *exsA* knockout mutant PAO1(Δ*exsA*). We constructed a single-copy chromosomally integrated CTX-*femA-lux* reporter in PAO1 and PAO1(Δ*exsA*) and compared the transcriptional activities by measuring the luminescence in these strains. In iron-limited M9 media supplemented with 1µM mycobactin J, the maximal promoter activity of *femA* was two-fold lower in PAO1(Δ*exsA*) compared with that in the wild-type PAO1 (Fig. 3A). The *femA* expression was undetectable in iron rich media (LB medium) (data not shown). The results indicate that the T3SS regulator ExsA plays a regulatory role in the expression of the *femA* operon, establishing a previously unknown link between the virulence factor T3SS and the Fem system in *P. aeruginosa*.

**FIG. 3.**
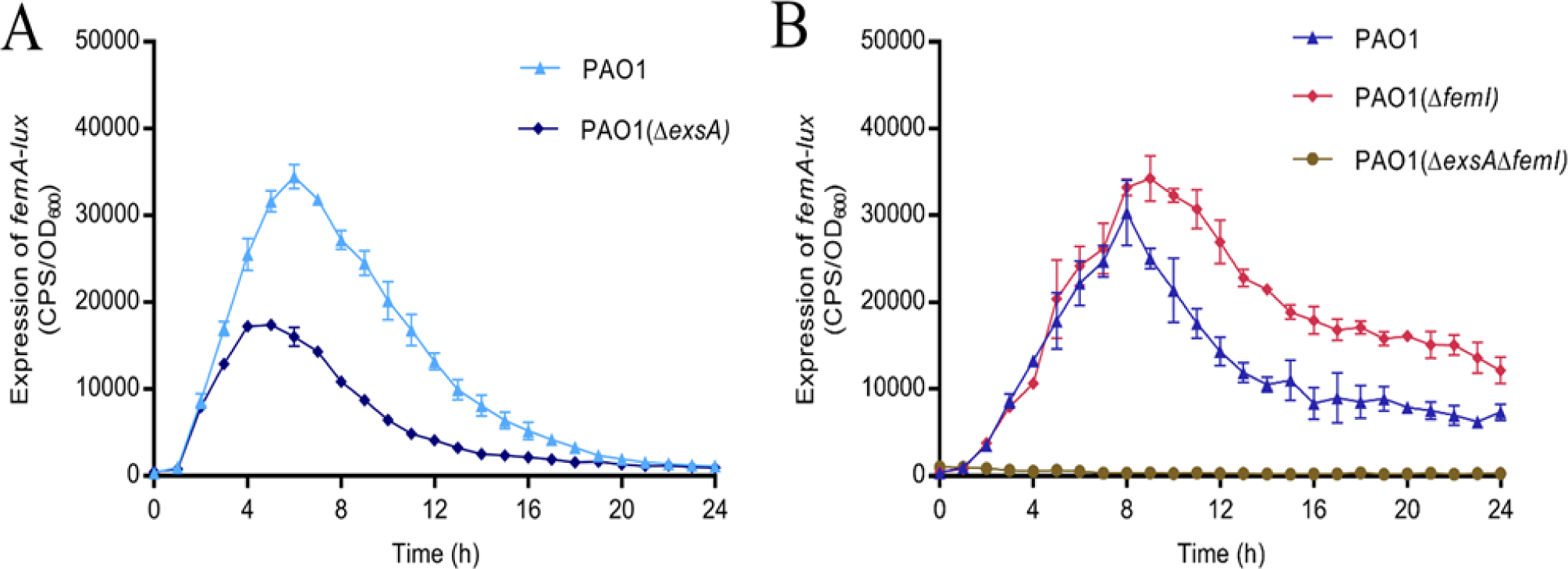
The regulation of the Fem system by both FemI and ExsA. A. Reduction of *femA* transcription levels in PAO1(Δ*exsA*) mutant compared with the wild-type PAO1. A CTX-*femA* reporter fusion was integrated into the chromosome of PAO1(Δ*exsA*) mutant and wild-type PAO1. *femA* promoter activity (solid lines) in the presence of mycobactin J (1 µM) was assessed over time. Bacterial growth (dotted lines) was monitored simultaneously (OD_600_). B. Comparison of *femA* expression in PAO1, PAO1(Δ*femI*) and PAO1(*ΔexsA*Δ*femI*). Slightly increased promoter activity of *femA* was observed in PAO1(Δ*femI*) but no activity was detected in PAO1(*ΔexsA*Δ*femI*). Promoter activity was recorded as (counts per second, cps) over 24 h every 30 minutes, and bacterial growth was monitored at the same time (OD_600_). The data represents an average of three independent experiments and error bars indicate standard deviations.

To clarify the relationship between this regulator ExsA and the sigma factor FemI and to determine whether ExsA activates *femA* expression in the absence of FemI, we constructed a *femI* deletion mutant, PAO1(Δ*femI*), and an *exsA* and *femI* double knock-out mutant, PAO1(Δ*exsA*Δ*femI*), and then we inserted the CTX-*femA* reporter into the chromosomes of PAO1(Δ*femI*) and PAO1(Δ*exsA*Δ*femI*). In iron limited M9 media supplemented with 1µM mycobactin J, the promoter activity of *femA* in PAO1(Δ*femI*) was evident despite of the lack of *femI* (Fig. 3B) and the expression level was actually slightly higher than in the wild type. However, the promoter activity of *femA* was not detectable in the double mutant PAO1(Δ*exsA*Δ*femI*) in the presence of mycobactin J (Fig. 3B). These results indicate that either ExsA or FemI was able to maintain *femA* expression in the absence of the other, but *femA* expression was abolished once both regulators were deleted. Clearly, in addition to FemI, ExsA can positively regulate *femA* expression, confirming a role of the T3SS regulator in the Fem surface signaling system.

### Fem system functions beyond its role in iron uptake

The reported function of the Fem system is the uptake of iron chelators (Mycobactin and Carboxymycobactin) produced by *Mycobacterium* species under iron-restricted conditions ^15^; hence, the usefulness of such a system in *P. aeruginosa* and its emergence in evolution would be hypothetically linked to its ability of scavenging siderophores produced by other bacteria in the vicinity. Given that *P. aeruginosa* synthesizes two potent siderophores, pyoverdine and pyochelin autonomously, one could reasonably speculate that the Fem system might have functions beyond mycobactin uptake. To examine such a possibility, we tested the effect of the presence of mycobactin on a set of *P. aeruginosa* virulence factors and genetic regulators. We compared the expression of these genes in the presence and absence of mycobactin J. As shown in Fig. 4A, among the genes tested, the expression of T3SS exotoxin gene *exoS* and *pilG* operon that encodes a component of type IV pili, was significantly increased in the presence of mycobactin J. The genes/operons with significantly decreased expression in the presence of mycobactin J include: *mexAB-oprM*, encoding a drug efflux pump of the resistance-nodulation-cell division (RND) family; *phzA1* and *phzA2* operons responsible for phenazine biosynthesis; *H2-T6SS* and *H3-T6SS,* which encode type VI protein secretion systems; *lasR* and *rhlR* of the quorum-sensing systems; *rhlA* responsible for the synthesis of rhamnolipids; and *rsmZ* encoding a small regulatory RNA (Fig. 4A). No significant changes were observed with the promoter activities of *xcpR*, *rpoS*, and *rnr*, regardless the presence of mycobactin J (data not shown). These results suggest that, in addition to the autoregulation of *femA* and *femI-femR*, the activation of Fem system by the presence of mycobactin J could affect key virulence factors in *P. aeruginosa*.

**FIG. 4.**
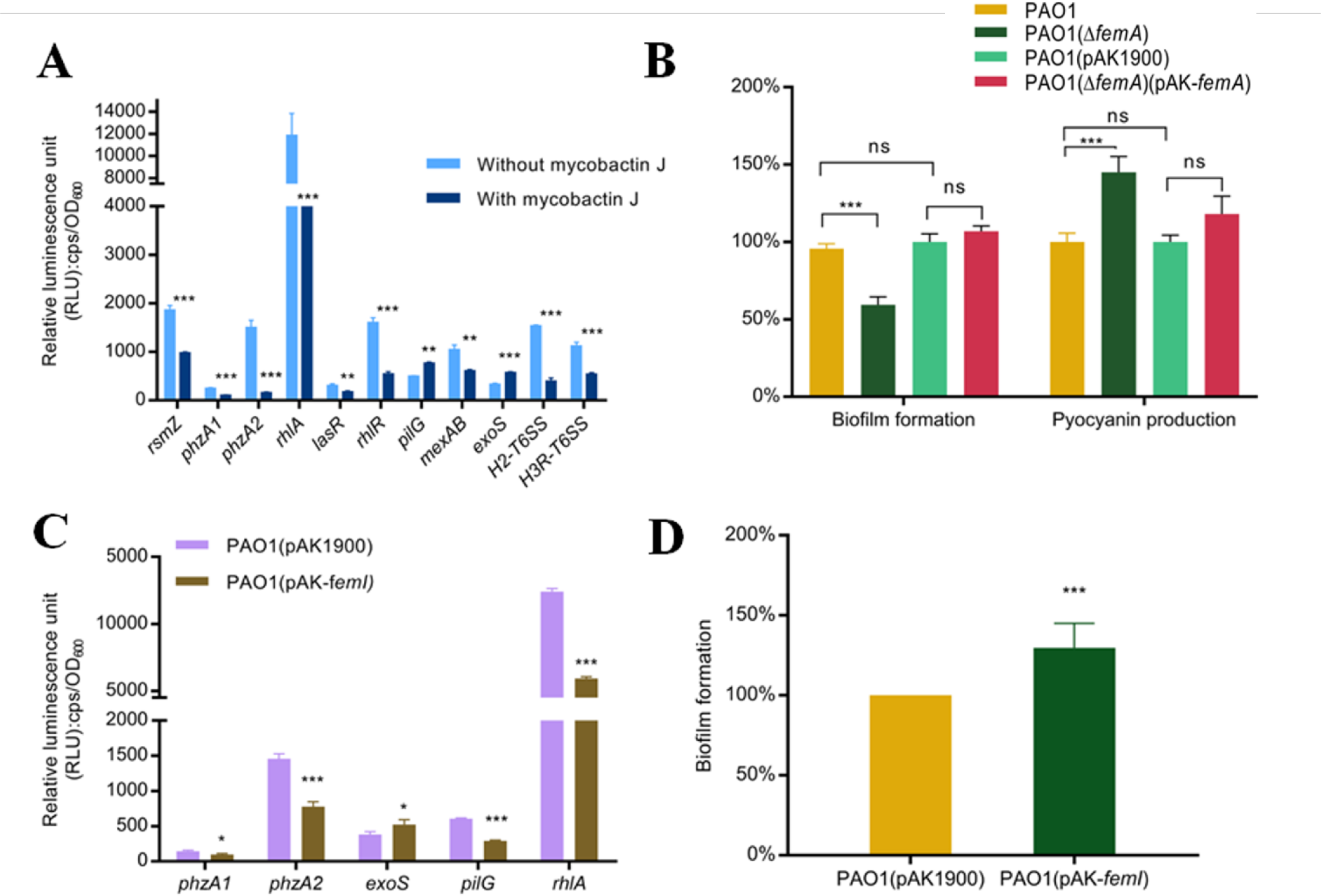
The effect of mycobactin J and the Fem system on the expression of virulence factors in *P. aeruginosa*. A. Differences in transcriptional activities of virulence factors and regulatory genes in the presence versus absence of mycobactin J. The peak promoter activities of the virulence genes were compared and presented as luminescence normalized to bacterial growth (RLU: cps/OD_600_). Unpaired Student’s t-test was used to analyze the data. Error bars indicate standard deviations. * (*p* < 0.05); **(*p* < 0.01); *** (*p* < 0.001). B. Effect of *femA* mutation on pyocyanin production and biofilm formation. The wild-type PAO1 was used as control and the data are presented as percent change relative to wild-type PAO1. These experiments were repeated at least three times. C. The effect of sigma factor FemI on virulence factor. D. Effect of *femI* overexpression on biofilm formation. The data is presented as percent change relative to wild-type PAO1.

Since iron is essential for bacterial survival and iron availability can serve as a signal for *P. aeruginosa* to regulate its virulence factors ^18^, it is possible the above-observed effect of Fem on those genes in the presence of mycobactin was merely due to the change of iron availability. To examine the potential existence of a potential regulatory effect by Fem itself, we constructed a *femA* knockout strain PAO1(Δ*femA*), and with this mutant we tested two virulence-related phenotypes that are connected to the above-mentioned genes (Fig. 4A) without the presence of mycobactin. As shown in Fig. 4B, without a change of iron availability, the production of pyocyanin was significantly increased in the PAO1(Δ*femA*), while biofilm formation was reduced significantly. Expression of *femA* gene on a plasmid introduced in the mutant strain PAO1(Δ*femA*) restored the pyocyanin production and the biofilm formation to the wild-type level. The results indicate that the Fem system influences some virulence factors in *P. aeruginosa*.

We further examined the effect of the Fem system on *P. aeruginosa* virulence factors through overexpression of *femI*. The full-length *femI* gene was cloned into pAK1900 and transferred into PAO1 to construct a *femI* overexpression strain, in which the promoter activity of several virulence factor genes was then assessed. As shown in Fig. 4C, the promoter activity of *exoS* increased significantly in the *femI* overexpression strain while the expression of *phzA1*, *phzA2*, *pilG* and *rhlA* decreased significantly. Furthermore, the *femI* expression strain produced significantly more biofilm than the control (Fig. 4D). These results indicate that the Fem system is able to affect virulence factors in *P. aeruginosa,* independent of its iron transportation role or iron availability in the experimental condition.

To further elucidate the roles of the Fem system, we carried out two sets of RNA-Seq analyses. First, we performed an RNA-Seq analysis to compare the mRNAs in PAO1 in the presence and absence of mycobactin J. Total RNA was extracted from cultures grown at mid-exponential phase, cDNA was synthesized to construct the cDNA library. Sequencing was performed using the Illumina HiSeq™ 2500 platform with pair-end 150 base reads.

As shown in Fig. 5, the expression of 113 genes in PAO1 was upregulated at least 2-fold in the presence of mycobactin J while the expression of 189 other genes was reduced 2-fold or more. The detailed information of these altered genes is listed in Table S3. Clearly, the uptake of mycobactin through the Fem system significantly affected the expression of large number of genes in *P. aeruginosa*. The RNA-Seq data corelated well with the results obtained using individual *lux*-based gene reporter fusions. The differentially expressed genes can be assigned to 27 functional groups. The GO categories significantly enriched (*q*< 0.05) among the differentially expressed genes are shown in Fig. 5A.

**FIG. 5.**
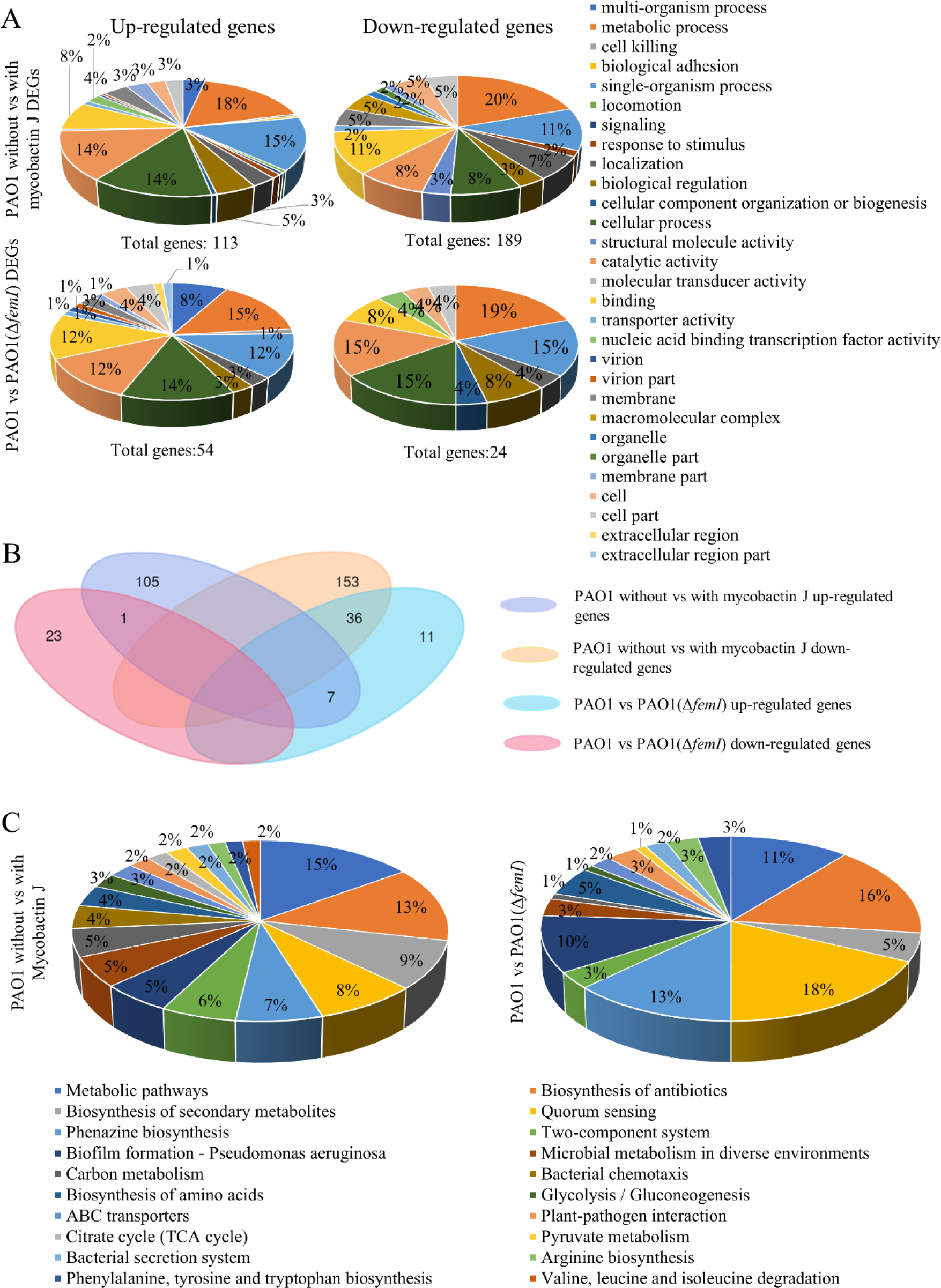
Gene expression profiling by RNA-Seq analysis. A. Pie charts showing gene ontology analysis results for the differentially expressed genes outlined in PAO1 without vs with mycobactin J, and PAO1 vs PAO1(Δ*femI*) in the presence of mycobactin J, respectively, subdivided into up-regulated (left) and down-regulated (right) groups. The abundance of each category is indicated as a percentage as well as the total number of genes included in each group. B. Venn diagram denoting the number of DEGs between PAO1 without vs with mycobactin and between PAO1(Δ*femI*) vs PAO1 in the presence of mycobactin J. C. Pie charts representing the distribution of top 20 KEGG pathways of differentially expressed genes in the two sets of RNA-Seq data.

To further study the role of Fem system in addition to mycobactin uptake, a second set of RNA-Seq was carried out to determine the regulon of FemI. We compared the transcriptional profiles of PAO1 and PAO1(Δ*femI*) in the presence of mycobactin J. The results indicate that 54 genes were upregulated in PAO1(Δ*femI*) while 24 genes were downregulated in the mutant. The GO categories that were significantly enriched (q< 0.05) are shown in Fig. 5A. These results demonstrate that the FemI activity not only involved in regulation of the *femA* operon but also affected other genes in *P. aeruginosa*. Comparing the genes affected by the presence of mycobactin J and those by the *femI* deletion, we uncovered several common and unique genes affected (Fig. 5B).

Analysing the expression patterns between PAO1 in the absence and presence of mycobactin J in terms of metabolic pathways, we found that the genes with altered expression included those participating in phenazine biosynthesis pathways, quorum sensing, biosynthesis of antibiotics, bacterial chemotaxis, host-pathogen interaction, and biofilm formation (Fig. 5C; Table S3). It is interesting to note that the genes involved in bacterial interspecies competition such as T6SS and antibiotic phenazine compound synthesis were downregulated in the presence of mycobactin J. Similarly, when *femI* is deleted, the genes affected also include those involved in phenazine biosynthesis, quorum sensing, biosynthesis of antibiotics, and biofilm formation (Fig. 5C; Table S4). *femI* deletion significantly decreased the expression of *rsmA* and increased the two pyocyanin synthesis operons, *phzA1* and *phzA2*. In addition, quorum sensing PQS system was highly expressed in the mutant. It is notable that two transcriptional regulators, MvaT and a putative LysR family transcriptional regulator PA3895, were significantly decreased in PAO1(Δ*femI*) while the levels of two other transcriptional regulators, VqsM and a hypothetical GntR regulator PA2299, were increased.

Altogether, it is evident that the FemI not only regulates *femA* and mycobactin-mediated iron uptake but can also affects multiple genes and activities in *P. aeruginosa*.

### The Fem system affects the virulence of P. aeruginosa in vivo

To verify the effect of the Fem system on *P. aeruginosa* pathogenicity *in vivo*, we used a *Galleria mellonella* infection model to examine the pathogenicity of different *P. aeruginosa* strains. The relative survival rates of the infected *G. mellonella* larvae were compared. As shown in Fig. 6, PAO1(Δ*femA*) exhibited significantly decreased mortality rate as compared to the wild-type PAO1. In agreement with this, *femI* overexpression strains displayed significantly higher mortality rate as compared with the control strain PAO1(pAK1900) (Fig. 6). The results show that the Fem system plays an important role in the *in vivo* pathogenicity of *P. aeruginosa*. Even in the absence of mycobactin, the Fem system seems to be required for the full virulence of *P. aeruginosa*.

**FIG. 6.**
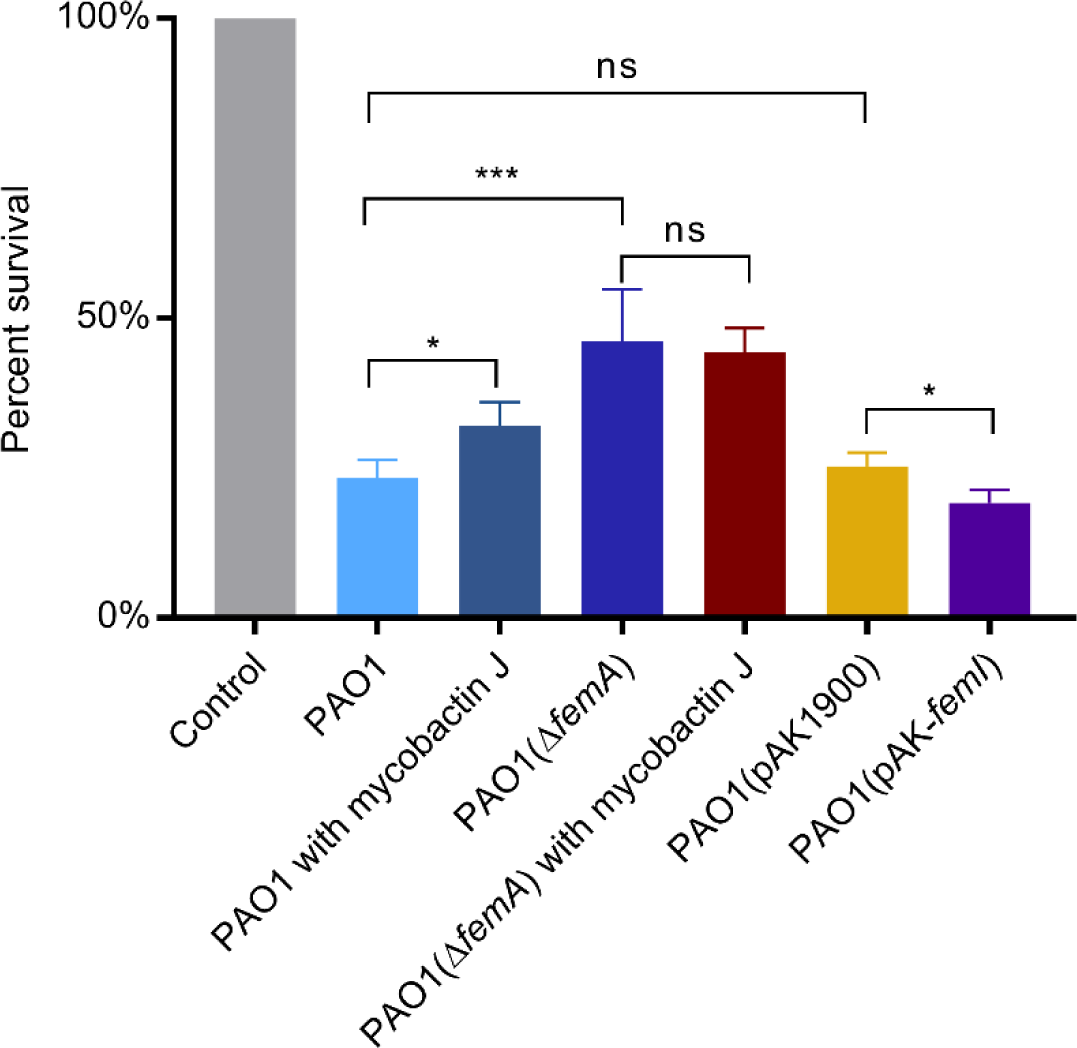
The effect of Fem system and mycobactin on the pathogenicity of *P. aeruginosa* shown in *G. mellonella* larva infection model. Larvae infected with different *P. aeruginosa* strains were maintained at 30 °C. The control group was inoculated with PBS to monitor for killing due to physical trauma. 18 h post infection, larvae were examined, and survival rates were calculated. The experiment was performed three times on different days, and unpaired Student’s t-test was used to analyze the data. Error bars indicate standard deviations. ***, *p* < 0.001, *, *p* <0.05, ns, not significant.

We further checked the effects of mycobactin J on the *in vivo* virulence. In the presence of mycobactin J, *P. aeruginosa* showed decreased pathogenicity in the *G. mellonella* infection model. However, no difference was observed in the *femA* deletion mutant regardless the presence of mycobactin J (Fig. 6), agreeing that mycobactin J’s effect on *P. aeruginosa* pathogenicity operates through the Fem system. The decreased pathogenicity of the wild-type *P. aeruginosa* in the presence of mycobactin J is also in agreement with the decreased expression of many other virulence factors in the presence of mycobactin J (Fig. 4).

## DISCUSSION

The Fem system in *P. aeruginosa* is believed to be an iron-transporting system that takes up iron-bound mycobactin produced by *Mycobacterium* spp. ^15^. The outer membrane protein FemA takes up mycobactin, which generates and transmits a signal to the anti-sigma factor FemR, releasing the sigma factor FemI to autoregulate *femI-femR* operon and *femA* operon. The regulation of the Fem system by the T3SS regulator ExsA revealed in this study is somewhat unexpected, considering that T3SS is a system used mainly for interacting with eukaryotic host cells. Our data indicate that the operon of *femA*-PA1909 bears typical ExsA-binding consensus sequences and the binding of ExsA to the promoter region was confirmed by EMSA. The expression of *femA* decreased significantly in the *exsA* knockout mutant PAO1(Δ*exsA*) under iron-deprived conditions. The deletion of *femI* only affected *femA* expression slightly with a small increase of expression observed in the *femI* mutant. However, no *femA* was expressed once *exsA* was also deleted, indicating that, without the presence of *femI*, *exsA* would be required for *femA* expression. These results showed that the T3SS regulator ExsA is able to regulate the expression of the Fem system, connecting the virulence factor with the Fem system for the first time.

Under iron-limited conditions, FemA serves as receptor to sense heterogenous siderophores (e.g. mycobactin) for *P. aeruginosa*. Interestingly, the results from the *G. mellonella* infection experiments indicate that the Fem system plays an important role in the pathogenicity of *P. aeruginosa*. In the presence of mycobactin J, the promoter activities of toxin secretion (*exoS)* and attachment factors (*pilG*) were increased, while the expression of other virulence factors including phenazine production, quorum-sensing, efflux pumps and T6SS secretion system (*phzA1*, *phzA2*, *rsmZ*, *rhlA*, *lasR*, *rhlR*, *pqs*, *mexAB, H2-T6SS, H3R-T6SS*) was significantly decreased. Such effects could be attributed partially to iron availability that was altered through mycobactin. As reported previously, genes responsible for rhamnolipid synthesis, *rhlA* and *rhlC*, as well as the genes of quorum sensing, are up-regulated in iron-limited conditions ^20^. H2-T6SS and H3-T6SS are also affected by iron availability ^21^. Therefore, the changed cellular iron level seems to explain some of the changes of gene expression observed in the presence of mycobactin.

However, not all the changes observed in our study could be explained by Fem system’s role in iron transport; the data suggests that the Fem system may play a role in regulating virulence factors in *P. aeruginosa,* independent of its iron transport function. First, the *P. aeruginosa femA* mutant, where a mycobactin effect was absent, produced significantly more pyocyanin than the wild-type PAO1 while it formed much less biofilm, indicating that the Fem system plays a role that is not connected to its mycobactin-transporting function. Second, the sigma factor FemI of the Fem system alone could affect virulence factors when expressed on a plasmid. Overexpression of *femI* decreased the promoter activities of *phzA1*, *phzA2*, *pilG* and *rhlA*, while it increased the promoter activities of *exoS.* Since no mycobactin was present in these experimental conditions, the effects of FemI on biofilm formation and the expression of virulence factors were independent of its iron acquisition function. The Fem system, when activated, affected gene expression and activities in *P. aeruginosa*. The RNA-Seq data also showed the deletion of *femI* significantly affected 78 genes when no mycobactin was involved. Lastly, the results from the animal study confirm that the Fem system plays an important role in the pathogenicity of *P. aeruginosa in vivo*, independent of its mycobactin-transporting function. Therefore, the Fem system appears to be integrated into the complex regulatory network that influences *P. aeruginosa* virulence. Further investigation is required to identify the exact mechanism of the Fem system’s involvement. However, the genes identified in the RNA-seq analysis provide a basis for future studies.

Several of the virulence factors affected by the Fem system are also involved in inter-species interactions. The Rhl system controls the synthesis of *P. aeruginosa* surfactants (rhamnolipids), which is involved in the killing of *S. aureus* by *P. aeruginosa* ^22^ and growth modulation of *Burkholderia cepacia* complex ^23^. Pyocyanin from *P. aeruginosa* is an antimicrobial agent against other competitors ^24^. Activation of such virulence factors by the Fem system may potentially benefit *P. aeruginosa* in inter-species competition ^25^. Mycobactin is regarded as a growth factor that enhances the growth of *Mycobacterium* species ^15^. Taking up mycobactin by the Fem system could also provide *P. aeruginosa* a competitive edge over *Mycobacterium* species. On the other hand, one of the *P. aeruginosa* responses triggered by the mycobactin and the Fem system was the decreased expression of T6SS system, a system used by *P. aeruginosa* to attack competitor bacteria. Such a response in *P. aeruginosa* could also benefit *Mycobacterium* species, the producers of mycobactin. In addition, the decreased production of antibiotic phenazine compounds by *P. aeruginosa* also favor the mycobactin producers. Clearly the role of the FEM system in inter-species interaction warrants further investigation.

## MATERIALS AND METHODS

### Bacterial strains and culture conditions

The bacterial strains and plasmids used in this study are described in Table S1. *Escherichia coli* and *P. aeruginosa* PAO1 were routinely grown on Luria-Bertani (LB) agar plates or in LB broth at 37°C unless otherwise specified. For iron-restricted conditions, *P. aeruginosa* cells were cultured in liquid CAS medium ^15^ or M9 minimal media, supplemented with 400 µM of 2, 2′-bipyridyl (Sigma-Aldrich). Mycobactin J was obtained from Allied Monitor (USA). When required, antibiotics were used at the following final concentrations (µg ml^−1^): for *P. aeruginosa*, gentamicin (Gm) at 50 and tetracycline (Tc) at 70 in LB or 300 in *Pseudomonas* isolation agar (PIA), streptomycin (Sm) at 150, carbenicillin (Cb) at 250 and trimethoprim (Tmp) at 300 in LB; for *E. coli*, kanamycin (Km) at 50, ampicillin (Amp) at 100, Sm at 12.5, Tc at 15 and Gm at 15 in LB.

### ExsA protein expression and purification

To elucidate the regulation of *exsA*, the expression of the protein ExsA was carried out by using Champion^TM^ pET Expression system (Invitrogen). Firstly, *exsA* was cloned into pET SUMO vector and transformed into BL21 (DE3) One Shot Chemically Competent *E. coli* cell. Successful constructs were confirmed with PCR and sequencing. The right transformant was inoculated into 50 ml LB broth containing 50 µg ml^−1^ kanamycin and grown to OD_600_ of 0.5. The cultures were then induced with 1mM isopropyl-b-Dthiogalactopyranoside (IPTG) for an additional 5 h at room temperature. ExsA_His_-containing bacterial lysate was cleared by centrifugation and subjected to purification by HisPur Ni-NTA Resin (Thermo Scientific) as per the manufacturer’s instruction.

### Electrophoretic mobility shift assay (EMSA)

To determine the binding specificity, EMSA was carried out using the purified ExsA_His_ protein and the 6-FAM^TM^ (fluorescein)-labeled DNA fragment containing the promoter regions of *femA*, *exoT* and *norC*. These labeled probes were amplified by PCR and purified by using clean-up kit (Qiagen). EMSA was performed as follows. Reaction (20 µl) containing 6-FAM^TM-^*femA* promoter (20 nM), indicated concentrations of protein ExsA, and 10 µl of 2×binding buffer [20 mM Tris (pH 7.5), 200 mM KCl, 2 mM EDTA, 2 mM dithiothreitol, 10% glycerol and 200 mg ml^−1^ bovine serum albumin] were incubated for 20 min at 25 °C. The unlabeled fragment of *femA* was added to the labeled probe at a ratio of 10:1 as a specific competitor. 6-FAM^TM^-*exoT* promoter was used as a positive control, and the unlabeled fragment of *norC* was used (10-fold) as a nonspecific competitor. Samples were then immediately subjected to electrophoresis on a 6.5% native polyacrylamide glycine gel at 4 °C. The band shifts were detected and captured by using Fusion FX7 Vilber Lourmat Imaging machine (Montreal Biotech Inc.).

### Construction of promoter-reporter strains and gene expression detecting systems

The plasmid pMS402 containing a promoterless *luxCDABE* reporter gene cluster was used to construct promoter-*luxCDABE* reporter fusions as described previously ^6^. Firstly, the promoter region of *femA* was generated by PCR and inserted into the vector pMS402 carrying promoterless *luxCDABE*, and the promoter-*luxCDABE* reporter cassette was then isolated and ligated into CTX6.1. The construct was then transformed into *E. coli* SM10-λ *pir* and the *P. aeruginosa* reporter integration strain CTX-*femA* in PAO1 was obtained by bi-parental mating. Using the *lux*-based reporter, gene expression in liquid culture was examined in a Synergy H4 Multimode Microplate Reader (BioTek). In brief, bacteria were cultured overnight in LB followed by dilution into fresh medium to OD_600_ of 0.2 and cultivated for an additional 3 h before use as inoculants. The cultures were inoculated into 96-well white plates with transparent bottom in triplicates in a ratio of 5 µl of inoculum to 95 µl of fresh medium. 50 µl of filter-sterilized mineral oil (Sigma Aldrich) was added on top to prevent evaporation during the assay. Luminescence (counts per second, cps) was measured every 30 min for 24 h. Bacterial growth was monitored at the same time by measuring OD_600_.

### Gene knockout strain construction

The knockout mutant was constructed by allelic exchange with the pEX18Tc sucrose counter-selection system as described previously ^26^. Briefly, the 0.5 kb upstream fragment of *femA* was amplified by PCR (the primers used are listed in Table S2) and firstly inserted into pEX18Tc resulting in pEX18Tc-*femA*_up_. Then, the 0.5 kb downstream fragment of *femA* was generated by PCR and ligated into pEX18Tc-*femA*_up_ to construct pEX18Tc-*femA*.The BamHI-digested Sm^r^-Ω cassette derived from pHP45Ω was inserted into the two PCR fragments of *femA* to generate pEX18Tc-*femA*-Ω. For unmarked deletion of *femI*, Gibson Assembly^®^ Cloning Kit was employed (New England BioLabs Inc) to make the deletion construct pEX-18Tc-*femI*. The upstream and downstream fragment of *femI* as well as the vector (pEX-18Tc) were amplified by PCR and directly added with the Gibson Assembly Master Mix, then incubated the reaction on a thermocycler at 50 °C for 15 minutes to assemble pEX-18Tc-*femI*. The *femA* knockout mutant, PAO1(Δ*femA*), was obtained by means of tri-parent mating as reported previously. Briefly, overnight cultures of the donor strain *E. coli* containing the plasmid pEX18Tc-*femA*-Ω, the helper strain containing pRK2013 and the recipient PAO1 were collected and re-suspended in PBS. The bacteria were mixed in a ratio of 2:2:1 and then spotted onto LB agar plates. After culturing at 37 °C overnight, the bacteria were scraped off and re-suspended in 500 µl LB. The diluted suspensions were spread on PIA plates containing streptomycin at 150 µg ml^−1^ and 10% sucrose. The resultant *femA* knockout mutant was verified by PCR and designated as PAO1(Δ*femA*). PAO1(Δ*femI*) and PAO1(Δ*exsA*Δ*femI*) were generated using the same strategy in PAO1 and PAO1(Δ*exsA*) background, respectively. These resultant mutants were verified by PCR and sequencing.

### RNA isolation

Strains were grown overnight at 37 °C followed by sub-culturing into fresh medium and grown to mid-exponential phase. Total RNA was extracted by TRIzol-based method (Life Technologies, CA, USA). In brief, the cultures were centrifuged at 12,000 x g for 5 minutes. The cell pellets were resuspended in 1 ml of TRIzol and then incubated at room temperature for 5 minutes to permit complete dissociation of nucleoproteins complex. After adding 0.2 ml of chloroform into the tube and vertex, the samples were centrifuged (12,000 x g at 4 °C for 15 minutes). The upper aqueous phase was transferred to a fresh tube and 0.5 ml of cold isopropanol was added to precipitate RNA. After centrifugation (12,000 x g for 10 min), the pellet was resuspended in 1 ml 75% ethanol and centrifuged again. The air-dried RNA pellet was finally dissolved in 50 µL RNase-free water.

### RNA-Seq library construction and sequencing

RNA integrity was measured using Bioanalyzer 2100 (Agilent, Santa Clara, CA) and rRNA was removed from 1 mg of total RNA with Ribo-Zero Magnetic Gold Kit (Epicentre Biotechnologies, Madison, WI, USA). To construct the RNA-Seq library, TruSeq RNA Sample Prep Kit v2 (Illumina, San Diego, CA, USA) was used. rRNA-depleted RNA was fragmented using Elute Prime Fragment Mix. First-strand cDNA was synthesized with First Strand Master Mix and Super Script II reverse transcriptase (Invitrogen, Carlsbad, CA, USA). Following purification by Agencourt RNAClean XP beads (Beckman Coulter, CA, USA), the second-strand cDNA library was synthesized using Second Strand Master Mix with dATP, dGTP, dCTP, dUTP mix. Purified fragmented cDNA was end repaired (30 minutes at 37 °C) prior to ligating sequencing adapters. Amplified RNA-Seq libraries were purified by using AMPureXP Beads. The clustering of the index-coded samples was performed on a cBot Cluster Generation System following to the manufacturer’s instructions, and the sequencing was performed using the Illumina Hiseq TM 2500 platform with pair-end 150 base reads.

### Bioinformatics analysis

The raw data from RNA-seq were filtered following three standards: 1, removing reads with ≥ 10 % unidentified nucleotides (N); 2, removing reads with > 50 % bases having Phred quality scores of ≤ 20; 3, removing reads aligned to the barcode adapter using FASTP (https://github.com/OpenGene/fastp). Quality trimmed reads were aligned using Bowtie2 (version 2.2.8) to the *P. aeruginosa* PAO1 reference genome to identify known genes and calculated gene expression by RSEM ^27^. The gene expression level was calculated and further normalized by using the fragments per kb of transcript per million (FPKM) mapped reads method to eliminate the influence of different gene lengths and amount of sequencing data on the calculation of gene expression. The edgeR package (http://www.r-project.org/) was used to identify differentially expressed genes (DEGs) across samples with fold changes ≥ 2 and with a false discovery rate-adjusted P (*q* value) < 0.05. Go terms and KEGG pathway were defined as being significantly enriched when the *q* value ≤ 0.05.

### Measurement of pyocyanin production

Pyocyanin was extracted from the culture supernatants after 18 h incubation and quantitated by a previously described protocol ^28^. Briefly, 3 ml of chloroform was added into 5 ml of the culture supernatant. After extraction, the chloroform layer was transferred to a new tube and mixed with 1 ml of 0.2 M HCl. Following centrifugation at 4,500 x g for 10 minutes, the top layer (0.2 M HCl) was removed, and its absorbance was measured at 520 nm. The concentrations obtained, expressed as micrograms of pyocyanin produced per milliliter of culture supernatant, were calculated using an extinction coefficient of 17.072 at 520 nm.

### Quantitation of biofilm formation

Biofilm production was quantified as previously described by O’Toole ^29^. Cells from overnight cultures were diluted at 1:100 in M63 minimal medium supplemented with magnesium sulfate, glucose and casamino acids, then inoculated into 96-well polystyrene microtiter plates (Costar) and grown at 37°C for 24 h. After incubation, the cultures were discarded, and the plate was gently submerged in a small tub of water to remove unattached cells and media components. A 125 µl of 0.1% crystal violet was added to each well and staining was allowed for 20 min at room temperature. Wells were rinsed three times with distilled water, and 125 µl of 30% acetic acid was added to dissolve the remaining crystal violet. A 100 µl portion of this solution was transferred to a new plate, and the absorbance was measured at 550 nm (OD_550_).

### Construction of femA and femI overexpression plasmids

The multi-copy-number *E. coli*-*P. aeruginosa* shuttle vector pAK1900 carrying a *lac* promoter ^30^ was used for complementation of *femA* in PAO1(Δ*femA*) and overexpression of the sigma factor (*femI*) in *P. aeruginosa*. The intact *femA* and *femI* genes were amplified by PCR from genomic DNA. The primers used are listed in Table S2. The PCR products were digested by BamHI and HindIII, and then ligated into vector pAK1900. The resultant plasmids were then transformed into *P. aeruginosa* by electroporation.

### Galleria mellonella *killing assays*

The *Galleria mellonella* infection model is a widely-accepted animal model and the experiments were performed as previously described ^31^. The larvae were stored in wood chips at 10 °C and used within two weeks from shipment. Prior to inoculation into *G. mellonella* caterpillars, *P. aeruginosa* cells were washed twice with PBS and then diluted in PBS to a final concentration of 1000 CFU ml^−1^ with or without mycobactin J as required. A 10 µl Hamilton syringe was used to inject 10 µl bacterial suspension into *G. mellonella* via the last left proleg. The infected larvae were incubated in a static incubator in the dark at 30 °C, the optimum temperature for insect growth and development. The number of dead caterpillars was scored after 18 hours post infection. Caterpillars were considered dead when they displayed no movement in response to touch.

### Data availability

The RNA-Seq data has been deposited in GeneBank under an accession number GSE164773.

## SUPLEMENTAL MATERIAL

Supplemental material is available onine.

## ACKNOWLEDGEMENTS

This study was supported by a grant from the Natural Sciences and Engineering Research Council of Canada (NSERC) (RGPIN-05864-2019) awarded to KD. The funders had no role in study design, data collection and interpretation, or the decision to submit the work for publication. We thank late Dr. Colin Dawes and the members of our lab for critical comments on the manuscript.

