## Supplemental Table 1-4 for "The Fem Cell-surface Signaling System is Regulated by ExsA in *Pseudomonas aeruginosa* and Affects Pathogenicity"

**Table S1.** Bacterial strains and plasmids used in this study.

| Bacterial strain or plasmid | Relevant characteristics/sequence | Source |
|---|---|---|
| ***E. coli* strains** | | |
| DH5α | $F^-$ φ80*lac*ZΔM15 Δ(*lacZYA-argF*) U169 *recA*1 *endA*1 *hsdR*17 ($r_k^-$, $m_k^+$) phoA supE44 $\lambda^-$ *thi*$^-$1 *gyrA*96 *relA*1 | Invitrogen |
| SM10-λ *pir* | Mobilizing strain, RP4 integrated in the chromosome; $Kn^r$ | [1] |
| BL21(DE3) | $F^-$ *ompT hsdS$_B$* ($r_B^-m_B^-$) *gal dcm* (DE3) | Invitrogen |
| Mach1™-T1$^R$ | $F^-$ Φ80*lac*ZΔM15 Δ*lac*X74 *hsd*R($r_K^-$ $m_K^+$) Δ*rec*A1398 *end*A1 *ton*A | Invitrogen |
| ***P. aeruginosa* strains** | | |
| PAO1 | Wild type, lab strain | [2] |
| PAO1(Δ*femA*) | *femA* knockout mutant of PAO1; $Sm^r$ | This study |
| PAO1(Δ*femI*) | *femI* knockout mutant of PAO1 | This study |
| PAO1(Δ*exsA*Δ*femI*) | *exsA* and *femI* double knockout mutant of PAO1; $Gm^r$ | This study |
| PAO1(Δ*exsA*) | *exsA* knockout mutant of PAO1; *exsA* ::$Gm^r$ | [3] |
| **Plasmids** | | |
| pMS402 | Expression reporter plasmid carrying the promoterless *luxCDABE*; $Km^r$ $Tmp^r$ | [2] |
| CTX-6.1 | Integration plasmid origins of plasmid mini-CTX-*lux*; $Tc^r$ | [3] |
| pET SUMO | Bacterial protein expression vector containing ubiquitin-like modifier (SUMO); $Km^r$ | Invitrogen |
| pHP45Ω | The pHP45 vector included a circle and the Ω fragment in linear form; Sm/$Spe^r$ | [4] |
| pRK2013 | Broad-host-range helper vector; $Tra^+$, $Km^r$ | [3] |
| pEX18Tc | *oriT*$^+$ *sacB*$^+$ gene replacement vector with multiple-cloning site from pUC18; $Tc^r$ | [5] |

| | | |
|---|---|---|
| pAK1900 | *E. coli-P. aeruginosa* shuttle cloning vector, Amp$^r$ | 3 |
| pAK-*femI* | pAK1900 with a 1024 bp fragment of PA1912-11 between BamHI and HindIII; Amp$^r$, Cb$^r$ | This study |
| pEX18Tc-*femA*$_{up}$ | pEX18Tc carrying the upstream fragment of *femA* | This study |
| pEX18Tc-*femA* | pEX18Tc carrying the upstream and downstream fragment of *femA* | This study |
| pEX18Tc-*femA*-Ω | pEX18Tc carrying the upstream, downstream fragment of *femA* and Ω-Sm$^r$ cassette, Sm$^r$ | This study |
| CTX-*femA* | Integration plasmid, CTX6.1 with a fragment of pKD-*femA* containing *femA* promoter region and *luxCDABE* gene; Km$^r$, Tmp$^r$, Tc$^r$ | This study |
| CTX-*exoS* | Integration plasmid, CTX6.1 with a fragment of pKD-*exoS* containing *exoS* promoter region and *luxCDABE* gene; Km$^r$, Tmp$^r$, Tc$^r$ | 3 |
| CTX-*rhlA* | Integration plasmid, CTX6.1 with a fragment of pKD-*rhlA* containing *rhlA* promoter region and *luxCDABE* gene; Km$^r$, Tmp$^r$, Tc$^r$ | 6 |
| CTX-*algD* | Integration plasmid, CTX6.1 with a fragment of pKD-*algD* containing *algD* promoter region and *luxCDABE* gene; Km$^r$, Tmp$^r$, Tc$^r$ | 6 |
| CTX-*phzA1* | Integration plasmid, CTX6.1 with a fragment of pKD-*phzA1* containing *phzA1* promoter region and *luxCDABE* gene; Km$^r$, Tmp$^r$, Tc$^r$ | 7 |
| CTX-*phzA2* | Integration plasmid, CTX6.1 with a fragment of pKD-*phzA2* containing *phzA2* promoter region and *luxCDABE* gene; Km$^r$, Tmp$^r$, Tc$^r$ | 7 |
| CTX-*pilG* | Integration plasmid, CTX6.1 with a fragment of pKD-*pilG* containing *pilG* promoter region and *luxCDABE* gene; Km$^r$, Tmp$^r$, Tc$^r$ | 6 |

| | | |
|---|---|---|
| CTX-*rsmA* | Integration plasmid, CTX6.1 with a fragment of pKD-*rsmA* containing *rsmA* promoter region and *luxCDABE* gene; $Km^r$, $Tmp^r$, $Tc^r$ | 3 |
| CTX-*rsmY* | Integration plasmid, CTX6.1 with a fragment of pKD-*rsmY* containing *rsmY* promoter region and *luxCDABE* gene; $Km^r$, $Tmp^r$, $Tc^r$ | 8 |
| CTX-*rsmZ* | Integration plasmid, CTX6.1 with a fragment of pKD-*rsmZ* containing *rsmZ* promoter region and *luxCDABE* gene; $Km^r$, $Tmp^r$, $Tc^r$ | 8 |
| CTX-*rhlR* | Integration plasmid, CTX6.1 with a fragment of pKD-*rhlR* containing *rhlR* promoter region and *luxCDABE* gene; $Km^r$, $Tmp^r$, $Tc^r$ | 6 |
| CTX-*mexAB* | Integration plasmid, CTX6.1 with a fragment of pKD-*MexAB* containing *MexAB* promoter region and *luxCDABE* gene; $Km^r$, $Tmp^r$, $Tc^r$ | 9 |

**Table S2.** Primers used in this study.

| Primer | Sequence (5'→3') † | Restriction site |
|---|---|---|
| *femA*-up-S | CGTGAATTCGCCTGAAGTACGAAGCCATC | EcoRI |
| *femA*-up-AS | TATGGATCCGTGATCAGCGAGGATTGCAG | BamHI |
| *femA*-dw-S | AATGGATCCGCACTGGGCAAGGACTACTG | BamHI |
| *femA*-dw-AS | GCCAAGCTTCCAGGAGAGCAGGTCGTAGA | HindIII |
| *femA*-promoter-S | ATACTCGAGCTACCGTCGCGGCGTACT | XhoI |
| *femA*-promoter-AS | TTAGGATCCGGGGCGATGTCATAGCTG | BamHI |
| *femI*-up-S | GTCGACCTGCAACAGCAACCTGGACCTGA | |
| *femI*-up-AS | TCCAGGCACTTAGAGATGGCTCACAGCGTC | |
| *femI*-dw-S | GCCATCTCTAAGTGCCTGGAGCTGATGGAA | |
| *femI*-dw-AS | TTGCATGCCTTATCGACGAAGATCTCGCCG | |
| *femI*-vector-L | GGTTGCTGTTGCAGGTCGACTCTAGAGGATC | |
| *femI*-vector-R | TTCGTCGATAAGGCATGCAAGCTTGGCACT | |
| pAK-*femI*-S | TATGGATCCCTGATCGGCGAAGAAAGGCT | BamHI |
| pAK-*femI*-AS | GTAAAGCTTGTAGAGATGGCTCACAGCGT | HindIII |
| pAK-*femA*-S | TAAGGATCCAGGCGTTGCCATCGACATAC | BamHI |
| pAK-*femA*-AS | CATAAGCTTGCGGTCATTGGCTGGTCATC | HindIII |
| *femI*-promoter-S | TATCTCGAGTCGATGGACGCCCTCAGCAG | XhoI |
| *femI*-promoter-AS | TTAGGATCCAGATGGCTCACAGCGTCGT | BamHI |
| EMSA-*femA*-S | CCGCCATCCCCGTGGCAACCGC | |
| EMSA-*femA*-AS | ACGGGCATGTACAGGGTTCCTC | |
| EMSA-*exoT*-S | ATATCCCATCGGGTTCTCCGCC | |
| EMSA-*exoT*-AS | CGGGTTCTGCTGAGATGATTGA | |

| | |
|---|---|
| EMSA-*norC*-S | TGAATAACTGAGTAAATCGTTCG |
| EMSA-*norC*-AS | TCTCGGTGTGGTAGGTCAGGGCC |

Note: [†], underlined are restriction site sequences.

**Table S3.** Differentially expressed genes in PAO1 in the presence and absence of mycobactin J.

| Gene ID | Gene name | Gene product | log2(Fold change) | P-value |
|---|---|---|---|---|
| **Down-regulated genes** | | | | |
| PA0007 | PA0007 | hypothetical protein | -1.216 | 0.000 |
| PA0105 | *coxB* | cytochrome c oxidase, subunit II | -2.117 | 0.000 |
| PA0106 | *coxA* | cytochrome c oxidase, subunit I | -2.880 | 0.000 |
| PA0122 | *rahU* | RahU | -1.457 | 0.000 |
| PA0156 | *triA* | Resistance-Nodulation-Cell Division (RND) triclosan efflux membrane fusion protein, TriA | -1.223 | 0.003 |
| PA0175 | *cheR2* | CheR2 | -1.512 | 0.001 |
| PA0176 | *aer2* | aerotaxis transducer Aer2 | -1.382 | 0.000 |
| PA0178 | PA0178 | probable two-component sensor | -1.984 | 0.000 |
| PA0179 | PA0179 | probable two-component response regulator | -1.511 | 0.000 |
| PA0180 | *cttP* | chemotactic transducer for trichloroethylene [positive chemotaxis], CttP | -1.281 | 0.000 |
| PA0211 | *mdcD* | malonate decarboxylase beta subunit | -3.123 | 0.000 |
| PA0263 | *hcpC* | secreted protein Hcp | -3.222 | 0.000 |
| PA0306.1 | PA0306.1 | Uncharacterized protein | -1.077 | 0.001 |
| PA0332 | PA0332 | hypothetical protein | -2.210 | 0.002 |
| PA0355 | *pfpI* | protease PfpI | -1.196 | 0.001 |
| PA0451 | PA0451 | conserved hypothetical protein | -2.103 | 0.000 |
| PA0468 | PA0468 | hypothetical protein | -1.551 | 0.002 |
| PA0477 | PA0477 | probable transcriptional regulator | -5.176 | 0.002 |
| PA0547 | PA0547 | probable transcriptional regulator | -1.322 | 0.000 |
| PA0575 | *rmcA* | redox regulator of c-di-GMP, RmcA | -1.033 | 0.001 |
| PA0586 | *ycgB* | conserved hypothetical protein | -1.107 | 0.000 |

| | | | | |
|---|---|---|---|---|
| PA0587 | *yeaH* | conserved hypothetical protein | -1.100 | 0.000 |
| PA0588 | *yeaG* | conserved hypothetical protein | -1.047 | 0.000 |
| PA0743 | PA0743 | probable 3-hydroxyisobutyrate dehydrogenase | -1.437 | 0.000 |
| PA0788 | PA0788 | hypothetical protein | -1.015 | 0.001 |
| PA0798 | *pmtA* | phospholipid methyltransferase | -1.446 | 0.002 |
| PA0837 | *slyD* | peptidyl-prolyl cis-trans isomerase SlyD | -1.124 | 0.000 |
| PA0852 | *cbpD* | chitin-binding protein CbpD precursor | -1.393 | 0.001 |
| PA0852.1 | PA0852.1 | Uncharacterized protein | -1.480 | 0.003 |
| PA0982 | PA0982 | hypothetical protein | -1.800 | 0.000 |
| PA0996 | *pqsA* | PqsA | -3.522 | 0.000 |
| PA0997 | *pqsB* | PqsB | -3.437 | 0.000 |
| PA0998 | *pqsC* | PqsC | -4.221 | 0.000 |
| PA0999 | *pqsD* | 3-oxoacyl-[acyl-carrier-protein] synthase III | -3.825 | 0.000 |
| PA1000 | *pqsE* | Quinolone signal response protein | -4.212 | 0.000 |
| PA1001 | *phnA* | anthranilate synthase component I | -4.900 | 0.000 |
| PA1002 | *phnB* | anthranilate synthase component II | -4.521 | 0.000 |
| PA1041 | PA1041 | probable outer membrane protein precursor | -1.713 | 0.000 |
| PA1051 | PA1051 | probable transporter | -1.655 | 0.000 |
| PA1053 | *slyB* | conserved hypothetical protein | -1.932 | 0.000 |
| PA1054 | *shaA* | ShaA | -1.385 | 0.001 |
| PA1190 | *yohC* | conserved hypothetical protein | -1.984 | 0.000 |
| PA1243 | PA1243 | probable sensor/response regulator hybrid | -1.131 | 0.002 |
| PA1316 | PA1316 | probable major facilitator superfamily (MFS) transporter | -1.154 | 0.001 |
| PA1323 | PA1323 | hypothetical protein | -1.323 | 0.000 |
| PA1358 | PA1358 | hypothetical protein | -3.877 | 0.000 |
| PA1463 | PA1463 | hypothetical protein | -1.400 | 0.001 |
| PA1474 | PA1474 | hypothetical protein | -1.024 | 0.001 |
| PA1481 | *ccmG* | cytochrome C biogenesis protein CcmG | -3.060 | 0.001 |
| PA1523 | *xdhB* | xanthine dehydrogenase | -1.212 | 0.000 |
| PA1529 | *lig* | DNA ligase | -1.079 | 0.002 |
| PA1656 | *hsiA2* | HsiA2 | -1.497 | 0.000 |

| | | | | |
|---|---|---|---|---|
| PA1657 | *hsiB2* | hsiB2 | -1.655 | 0.001 |
| PA1730 | PA1730 | conserved hypothetical protein | -1.049 | 0.001 |
| PA1732 | PA1732 | conserved hypothetical protein | -1.291 | 0.002 |
| PA1753 | PA1753 | conserved hypothetical protein | -1.286 | 0.000 |
| PA1761 | PA1761 | hypothetical protein | -1.569 | 0.001 |
| PA1828 | PA1828 | probable short-chain dehydrogenase | -1.730 | 0.001 |
| PA1860 | PA1860 | hypothetical protein | -1.583 | 0.000 |
| PA1871 | *lasA* | protease LasA | -1.995 | 0.001 |
| PA1880 | PA1880 | probable oxidoreductase | -1.084 | 0.000 |
| PA1899 | *phzA2* | probable phenazine biosynthesis protein | -14.518 | 0.002 |
| PA1900 | *phzB2* | probable phenazine biosynthesis protein | -2.632 | 0.000 |
| PA1901 | *phzC2* | phenazine biosynthesis protein PhzC | -1.635 | 0.000 |
| PA1903 | *phzE2* | phenazine biosynthesis protein PhzE | -2.733 | 0.000 |
| PA1904 | *phzF2* | probable phenazine biosynthesis protein | -3.184 | 0.000 |
| PA1946 | *rbsB* | binding protein component precursor of ABC ribose transporter | -1.270 | 0.001 |
| PA2068 | PA2068 | probable major facilitator superfamily (MFS) transporter | -2.709 | 0.002 |
| PA2069 | PA2069 | probable carbamoyl transferase | -1.714 | 0.000 |
| PA2153 | *glgB* | 1,4-alpha-glucan branching enzyme | -1.117 | 0.001 |
| PA2166 | PA2166 | hypothetical protein | -1.409 | 0.001 |
| PA2190 | PA2190 | conserved hypothetical protein | -1.567 | 0.000 |
| PA2247 | *bkdA1* | 2-oxoisovalerate dehydrogenase (alpha subunit) | -2.022 | 0.000 |
| PA2249 | *bkdB* | branched-chain alpha-keto acid dehydrogenase (lipoamide component) | -2.179 | 0.002 |
| PA2250 | *lpdV* | lipoamide dehydrogenase-Val | -2.861 | 0.000 |
| PA2291 | *oprB2* | probable glucose-sensitive porin | -1.287 | 0.000 |
| PA2293 | PA2293 | hypothetical protein | -3.200 | 0.000 |
| PA2294 | PA2294 | probable ATP-binding component of ABC transporter | -5.622 | 0.000 |
| PA2295 | PA2295 | probable permease of ABC transporter | -16.362 | 0.000 |
| PA2296 | PA2296 | hypothetical protein | -4.641 | 0.000 |

| | | | | |
|---|---|---|---|---|
| PA2297 | PA2297 | probable ferredoxin | -2.792 | 0.000 |
| PA2298 | PA2298 | probable oxidoreductase | -3.069 | 0.000 |
| PA2299 | PA2299 | probable transcriptional regulator | -2.644 | 0.000 |
| PA2300 | *chiC* | chitinase | -2.203 | 0.000 |
| PA2360 | *hsiA3* | hypothetical protein | -1.361 | 0.001 |
| PA2371 | *clpV3* | ClpV3 | -1.311 | 0.000 |
| PA2372 | PA2372 | hypothetical protein | -1.110 | 0.000 |
| PA2373 | *vgrG3* | VgrG3 | -1.358 | 0.000 |
| PA2443 | *sdaA* | L-serine dehydratase | -1.405 | 0.000 |
| PA2481 | PA2481 | hypothetical protein | -1.421 | 0.001 |
| PA2562 | PA2562 | hypothetical protein | -1.292 | 0.002 |
| PA2566 | PA2566 | conserved hypothetical protein | -1.517 | 0.000 |
| PA2570 | *lecA* | LecA | -14.867 | 0.002 |
| PA2572 | PA2572 | probable two-component response regulator | -1.017 | 0.001 |
| PA2573 | PA2573 | probable chemotaxis transducer | -1.082 | 0.000 |
| PA2621 | *clpS* | ATP-dependent Clp protease adaptor protein ClpS | -1.061 | 0.000 |
| PA2633 | PA2633 | hypothetical protein | -1.504 | 0.000 |
| PA2708 | PA2708 | hypothetical protein | -1.570 | 0.002 |
| PA2788 | PA2788 | probable chemotaxis transducer | -1.322 | 0.000 |
| PA2853 | *oprI* | Outer membrane lipoprotein OprI precursor | -1.470 | 0.000 |
| PA2920 | PA2920 | probable chemotaxis transducer | -1.613 | 0.000 |
| PA3023 | PA3023 | conserved hypothetical protein | -1.582 | 0.000 |
| PA3089 | PA3089 | hypothetical protein | -1.407 | 0.001 |
| PA3142 | PA3142 | integrase | -1.149 | 0.001 |
| PA3183 | *zwf* | glucose-6-phosphate 1-dehydrogenase | -1.521 | 0.000 |
| PA3185 | PA3185 | hypothetical protein | -1.009 | 0.002 |
| PA3194 | *edd* | phosphogluconate dehydratase | -1.090 | 0.000 |
| PA3195 | *gapA* | glyceraldehyde 3-phosphate dehydrogenase | -1.683 | 0.000 |
| PA3250 | PA3250 | hypothetical protein | -1.323 | 0.001 |
| PA3251 | PA3251 | hypothetical protein | -3.727 | 0.001 |
| PA3305 | PA3305 | hypothetical protein | -1.352 | 0.002 |
| PA3307 | PA3307 | hypothetical protein | -1.126 | 0.000 |

| | | | | |
|---|---|---|---|---|
| PA3354 | PA3354 | hypothetical protein | -1.547 | 0.003 |
| PA3356 | *pauA5* | glutamylpolyamine synthetase | -1.005 | 0.001 |
| PA3361 | *lecB* | fucose-binding lectin PA-IIL | -2.569 | 0.000 |
| PA3415 | PA3415 | probable dihydrolipoamide acetyltransferase | -1.414 | 0.000 |
| PA3416 | PA3416 | probable pyruvate dehydrogenase E1 component, beta chain | -1.667 | 0.000 |
| PA3417 | PA3417 | probable pyruvate dehydrogenase E1 component, alpha subunit | -1.144 | 0.001 |
| PA3418 | *ldh* | leucine dehydrogenase | -1.035 | 0.000 |
| PA3427 | PA3427 | probable short-chain dehydrogenases | -2.204 | 0.003 |
| PA3430 | PA3430 | probable aldolase | -1.477 | 0.000 |
| PA3431 | *ywbG* | conserved hypothetical protein | -2.409 | 0.000 |
| PA3432 | PA3432 | hypothetical protein | -2.136 | 0.000 |
| PA3479 | *rhlA* | rhamnosyltransferase chain A | -0.841 | 0.004 |
| PA3526 | *motY* | MotY | -1.101 | 0.003 |
| PA3582 | *glpK* | glycerol kinase | -1.965 | 0.000 |
| PA3584 | *glpD* | glycerol-3-phosphate dehydrogenase | -2.204 | 0.000 |
| PA3587 | *metR* | transcriptional regulator MetR | -1.382 | 0.000 |
| PA3628 | *yeiG* | putative esterase | -2.456 | 0.000 |
| PA3629 | *adhC* | alcohol dehydrogenase class III | -1.787 | 0.000 |
| PA3667 | PA3667 | probable pyridoxal-phosphate dependent enzyme | -1.220 | 0.001 |
| PA3684 | PA3684 | hypothetical protein | -2.066 | 0.000 |
| PA3688 | PA3688 | hypothetical protein | -1.372 | 0.002 |
| PA3691 | PA3691 | hypothetical protein | -1.501 | 0.000 |
| PA3692 | *lptF* | lipotoxon F, LptF | -1.209 | 0.000 |
| PA3723 | *yqjM* | probable FMN oxidoreductase | -1.251 | 0.000 |
| PA3808 | *yfhJ* | conserved hypothetical protein | -1.869 | 0.000 |
| PA3809 | *fdx2* | ferredoxin [2Fe-2S] | -1.405 | 0.000 |
| PA3846 | PA3846 | hypothetical protein | -1.467 | 0.000 |
| PA3851 | PA3851 | hypothetical protein | -1.609 | 0.001 |
| PA3852 | PA3852 | hypothetical protein | -1.201 | 0.002 |
| PA3878 | *narX* | two-component sensor NarX | -1.112 | 0.002 |

| | | | | |
|---|---|---|---|---|
| PA3920 | *copA1* | CopA1 | -1.708 | 0.000 |
| PA3921 | PA3921 | probable transcriptional regulator | -1.446 | 0.000 |
| PA3923 | PA3923 | hypothetical protein | -1.575 | 0.002 |
| PA3944 | PA3944 | conserved hypothetical protein | -1.317 | 0.003 |
| PA3969 | PA3969 | conserved hypothetical protein | -1.632 | 0.001 |
| PA4005 | *ybeB* | conserved hypothetical protein | -1.878 | 0.001 |
| PA4012 | PA4012 | hypothetical protein | -1.771 | 0.000 |
| PA4024 | *eutB* | ethanolamine-ammonia lyase, large subunit, EutB | -2.054 | 0.000 |
| PA4025 | *eutC* | ethanolamine-ammonia lyase, small subunit, EutC | -2.408 | 0.002 |
| PA4045 | *btuF* | conserved hypothetical protein | -3.401 | 0.000 |
| PA4048 | PA4048 | hypothetical protein | -4.681 | 0.000 |
| PA4141 | PA4141 | hypothetical protein | -2.038 | 0.000 |
| PA4142 | PA4142 | probable secretion protein | -2.860 | 0.000 |
| PA4143 | *cyaB* | probable toxin transporter | -2.537 | 0.000 |
| PA4144 | *opmK* | probable outer membrane protein precursor | -3.401 | 0.000 |
| PA4159 | *fepB* | ferrienterobactin-binding periplasmic protein precursor FepB | -1.310 | 0.000 |
| PA4214 | *phzE1* | phenazine biosynthesis protein PhzE | -2.733 | 0.000 |
| PA4215 | *phzF1* | probable phenazine biosynthesis protein | -3.184 | 0.000 |
| PA4216 | *phzG1* | probable pyridoxamine 5'-phosphate oxidase | -2.781 | 0.000 |
| PA4217 | *phzS* | flavin-containing monooxygenase | -2.744 | 0.000 |
| PA4421 | *yabB* | conserved hypothetical protein | -1.221 | 0.000 |
| PA4502 | *dppA4* | probable binding protein component of ABC transporter | -1.635 | 0.001 |
| PA4525 | *pilA* | type 4 fimbrial precursor PilA | -1.518 | 0.000 |
| PA4607 | PA4607 | hypothetical protein | -1.677 | 0.000 |
| PA4683 | PA4683 | hypothetical protein | -1.563 | 0.002 |
| PA4738 | PA4738 | conserved hypothetical protein | -1.393 | 0.000 |
| PA4739 | PA4739 | conserved hypothetical protein | -2.586 | 0.000 |
| PA4764 | *fur* | ferric uptake regulation protein | -1.137 | 0.002 |
| PA4781 | PA4781 | cyclic di-GMP phosphodiesterase | -1.124 | 0.002 |
| PA4880 | PA4880 | probable bacterioferritin | -1.600 | 0.000 |

| | | | | |
|---|---|---|---|---|
| PA4888 | desB | acyl-CoA delta-9-desaturase, DesB | -4.224 | 0.001 |
| PA4916 | *nrtR* | Nudix-related transcriptional regulator NrtR | -1.936 | 0.000 |
| PA4917 | *nadD2* | nicotinate mononucleotide adenylyltransferase NadD2 | -2.752 | 0.000 |
| PA4929 | PA4929 | hypothetical protein | -1.458 | 0.000 |
| PA4958 | *fimW* | FimW | -1.069 | 0.000 |
| PA5066 | *hisI* | phosphoribosyl-AMP cyclohydrolase | -1.424 | 0.000 |
| PA5105 | *hutC* | histidine utilization repressor HutC | -2.191 | 0.001 |
| PA5106 | PA5106 | conserved hypothetical protein | -5.705 | 0.000 |
| PA5107 | *blc* | outer membrane lipoprotein Blc | -1.334 | 0.002 |
| PA5161 | *rmlB* | dTDP-D-glucose 4,6-dehydratase | -1.055 | 0.000 |
| PA5171 | *arcA* | arginine deiminase | -1.644 | 0.000 |
| PA5172 | *arcB* | ornithine carbamoyltransferase, catabolic | -1.865 | 0.000 |
| PA5173 | *arcC* | carbamate kinase | -1.963 | 0.000 |
| PA5208 | PA5208 | conserved hypothetical protein | -1.434 | 0.003 |
| PA5271 | PA5271 | hypothetical protein | -1.479 | 0.001 |
| PA5303 | PA5303 | conserved hypothetical protein | -2.041 | 0.000 |
| PA5306 | *ynbE* | conserved hypothetical protein | -3.384 | 0.001 |
| **Up-regulated genes** | | | | |
| PA0094 | *eagT6* | EagT6 | 1.845 | 0.000 |
| PA0126 | PA0126 | hypothetical protein | 2.870 | 0.001 |
| PA0213 | PA0213 | hypothetical protein | 4.472 | 0.003 |
| PA0433 | PA0433 | hypothetical protein | 1.454 | 0.000 |
| PA0434 | PA0434 | hypothetical protein | 1.315 | 0.000 |
| PA0435 | PA0435 | hypothetical protein | 2.143 | 0.000 |
| PA0610 | *prtN* | transcriptional regulator PrtN | 2.285 | 0.000 |
| PA0612 | *ptrB* | repressor, PtrB | 3.626 | 0.000 |
| PA0613 | PA0613 | hypothetical protein | 2.478 | 0.000 |
| PA0614 | PA0614 | hypothetical protein | 2.641 | 0.000 |
| PA0615 | PA0615 | hypothetical protein | 1.929 | 0.000 |
| PA0616 | PA0616 | hypothetical protein | 2.798 | 0.000 |
| PA0617 | PA0617 | probable bacteriophage protein | 2.691 | 0.000 |

| | | | | |
|---|---|---|---|---|
| PA0618 | PA0618 | probable bacteriophage protein | 3.112 | 0.000 |
| PA0619 | PA0619 | probable bacteriophage protein | 2.787 | 0.000 |
| PA0620 | PA0620 | probable bacteriophage protein | 2.875 | 0.000 |
| PA0621 | PA0621 | conserved hypothetical protein | 3.222 | 0.000 |
| PA0622 | PA0622 | probable bacteriophage protein | 2.978 | 0.000 |
| PA0623 | PA0623 | probable bacteriophage protein | 3.242 | 0.000 |
| PA0624 | PA0624 | hypothetical protein | 2.989 | 0.000 |
| PA0625 | PA0625 | hypothetical protein | 3.006 | 0.000 |
| PA0626 | PA0626 | hypothetical protein | 3.144 | 0.000 |
| PA0627 | PA0627 | conserved hypothetical protein | 3.336 | 0.000 |
| PA0628 | PA0628 | conserved hypothetical protein | 2.858 | 0.000 |
| PA0629 | PA0629 | conserved hypothetical protein | 2.909 | 0.000 |
| PA0630 | PA0630 | hypothetical protein | 1.779 | 0.000 |
| PA0631 | PA0631 | hypothetical protein | 15.500 | 0.001 |
| PA0633 | PA0633 | hypothetical protein | 3.192 | 0.000 |
| PA0634 | PA0634 | hypothetical protein | 2.821 | 0.000 |
| PA0635 | PA0635 | hypothetical protein | 2.222 | 0.000 |
| PA0636 | PA0636 | hypothetical protein | 2.914 | 0.000 |
| PA0637 | PA0637 | conserved hypothetical protein | 2.757 | 0.000 |
| PA0638 | PA0638 | probable bacteriophage protein | 2.359 | 0.000 |
| PA0639 | PA0639 | conserved hypothetical protein | 2.932 | 0.000 |
| PA0640 | PA0640 | probable bacteriophage protein | 2.851 | 0.000 |
| PA0641 | PA0641 | probable bacteriophage protein | 2.895 | 0.000 |
| PA0642 | PA0642 | hypothetical protein | 2.848 | 0.000 |
| PA0643 | PA0643 | hypothetical protein | 3.033 | 0.000 |
| PA0644 | PA0644 | hypothetical protein | 3.071 | 0.000 |
| PA0645 | PA0645 | hypothetical protein | 3.084 | 0.000 |
| PA0646 | PA0646 | hypothetical protein | 2.716 | 0.000 |
| PA0647 | PA0647 | hypothetical protein | 2.697 | 0.000 |
| PA0648 | PA0648 | hypothetical protein | 2.277 | 0.000 |
| PA0649 | *trpG* | anthranilate synthase component II | 1.463 | 0.000 |
| PA0654 | *speD* | S-adenosylmethionine decarboxylase proenzyme | 2.446 | 0.000 |

| | | | | |
|---|---|---|---|---|
| PA0685 | *hxcQ* | HxcQ | 4.668 | 0.002 |
| PA0781 | PA0781 | hypothetical protein | 4.566 | 0.000 |
| PA0807 | *ampDh3* | AmpDh3 | 2.759 | 0.000 |
| PA0808 | PA0808 | hypothetical protein | 2.510 | 0.000 |
| PA0819 | PA0819 | hypothetical protein | 4.073 | 0.001 |
| PA0907 | *alpA* | lysis phenotype activator, AlpA | 2.135 | 0.000 |
| PA0908 | *alpC* | AlpC | 2.328 | 0.002 |
| PA0910 | *alpD* | AlpD | 2.247 | 0.000 |
| PA0911 | *alpE* | AlpE | 3.096 | 0.000 |
| PA0931 | *pirA* | ferric enterobactin receptor PirA | 1.136 | 0.001 |
| PA0985 | *pyoS5* | pyocin S5 | 3.454 | 0.000 |
| PA1150 | *pys2* | pyocin S2 | 2.088 | 0.000 |
| PA1152 | PA1152 | hypothetical protein | 4.726 | 0.000 |
| PA1168 | PA1168 | hypothetical protein | 3.583 | 0.000 |
| PA1169 | PA1169 | probable lipoxygenase | 3.534 | 0.000 |
| PA1374 | PA1374 | hypothetical protein | 4.449 | 0.003 |
| PA1596 | *htpG* | heat shock protein HtpG | 1.505 | 0.000 |
| PA1907 | PA1907 | hypothetical protein | 3.072 | 0.000 |
| PA1908 | PA1908 | probable major facilitator superfamily (MFS) transporter | 2.975 | 0.000 |
| PA1909 | PA1909 | hypothetical protein | 3.772 | 0.000 |
| PA1910 | *femA* | ferric-mycobactin receptor, FemA | 2.544 | 0.000 |
| PA1921 | PA1921 | hypothetical protein | 2.591 | 0.000 |
| PA1922 | *cirA* | probable TonB-dependent receptor | 4.221 | 0.000 |
| PA1923 | PA1923 | hypothetical protein | 6.367 | 0.000 |
| PA1925 | PA1925 | hypothetical protein | 6.129 | 0.000 |
| PA2381 | PA2381 | hypothetical protein | 1.140 | 0.000 |
| PA2393 | PA2393 | putative dipeptidase | 1.271 | 0.000 |
| PA2394 | *pvdN* | PvdN | 1.252 | 0.000 |
| PA2530 | PA2530 | hypothetical protein | 1.485 | 0.001 |
| PA2781 | PA2781 | hypothetical protein | 1.579 | 0.000 |
| PA2911 | PA2911 | probable TonB-dependent receptor | 1.776 | 0.000 |

| | | | | |
|---|---|---|---|---|
| PA3126 | *ibpA* | heat-shock protein IbpA | 1.556 | 0.000 |
| PA3413 | *yebG* | conserved hypothetical protein | 1.005 | 0.001 |
| PA3598 | *ypqQ* | conserved hypothetical protein | 2.973 | 0.000 |
| PA3600 | *rpl36* | conserved hypothetical protein | 2.755 | 0.000 |
| PA3601 | *ykgM* | conserved hypothetical protein | 2.581 | 0.000 |
| PA3656 | *rpsB* | 30S ribosomal protein S2 | 1.075 | 0.000 |
| PA3785 | PA3785 | conserved hypothetical protein | 1.923 | 0.000 |
| PA3866 | PA3866 | Pyocin S4 | 1.680 | 0.000 |
| PA4139 | PA4139 | hypothetical protein | 2.030 | 0.000 |
| PA4140 | PA4140 | hypothetical protein | 1.616 | 0.000 |
| PA4170 | PA4170 | hypothetical protein | 2.284 | 0.000 |
| PA4385 | *groEL* | GroEL protein | 1.064 | 0.000 |
| PA4431 | PA4431 | probable iron-sulfur protein | 1.334 | 0.000 |
| PA4530 | PA4530 | conserved hypothetical protein | 4.854 | 0.002 |
| PA4563 | *rpsT* | 30S ribosomal protein S20 | 1.245 | 0.002 |
| PA4625 | *cdrA* | cyclic diguanylate-regulated TPS partner A, CdrA | 1.031 | 0.003 |
| PA4640 | *mqoB* | malate:quinone oxidoreductase | 1.750 | 0.000 |
| PA4762 | *grpE* | heat shock protein GrpE | 1.488 | 0.000 |
| PA4772 | PA4772 | probable ferredoxin | 1.616 | 0.000 |
| PA4834 | *cntI* | CntI | 3.349 | 0.000 |
| PA4835 | *cntM* | CntM | 3.343 | 0.000 |
| PA4836 | *cntL* | CntL | 4.270 | 0.000 |
| PA4837 | *cntO* | CntO | 4.230 | 0.000 |
| PA4838 | PA4838 | hypothetical protein | 3.129 | 0.000 |
| PA5053 | *hslV* | heat shock protein HslV | 1.660 | 0.000 |
| PA5201 | *yhgF* | conserved hypothetical protein | 1.465 | 0.001 |
| PA5383 | *yeiH* | conserved hypothetical protein | 1.218 | 0.000 |
| PA5491 | PA5491 | probable cytochrome | 2.178 | 0.002 |
| PA5534 | PA5534 | hypothetical protein | 3.240 | 0.000 |
| PA5535 | PA5535 | conserved hypothetical protein | 2.869 | 0.000 |
| PA5536 | *dksA2* | DksA2 | 3.576 | 0.000 |
| PA5537 | PA5537 | hypothetical protein | 2.989 | 0.001 |

| | | | | |
|---|---|---|---|---|
| PA5538 | *amiA* | N-acetylmuramoyl-L-alanine amidase | 3.139 | 0.000 |
| PA5539 | PA5539 | RidA subfamily protein | 2.473 | 0.001 |
| PA5540 | PA5540 | hypothetical protein | 4.253 | 0.000 |
| PA5541 | *pyrQ* | dihydroorotase | 4.328 | 0.000 |
| PA5569 | *rnpA* | ribonuclease P protein component | 1.066 | 0.002 |

**Table S4.** Differentially expressed genes in PAO1 vs PAO1(Δ*femI*) in the presence of mycobactin J.

| **Gene ID** | **Gene name** | **Gene product** | **$\log_2$(Fold change)** | **P-value** |
|---|---|---|---|---|
| **Up-regulated genes** | | | | |
| PA0156 | *triA* | resistance-Nodulation-Cell Division (RND) triclosan efflux membrane fusion protein, TriA | 1.275 | 0.001 |
| PA0852 | *cbpD* | chitin-binding protein CbpD precursor | 1.653 | 0.000 |
| PA0852.1 | PA0852.1 | uncharacterized protein | 1.445 | 0.000 |
| PA0996 | *pqsA* | PqsA | 3.308 | 0.000 |
| PA0997 | *pqsB* | PqsB | 3.957 | 0.000 |
| PA0998 | *pqsC* | PqsC | 4.443 | 0.000 |
| PA0999 | *pqsD* | 3-oxoacyl-[acyl-carrier-protein] synthase III | 3.444 | 0.000 |
| PA1000 | *pqsE* | quinolone signal response protein | 4.126 | 0.000 |
| PA1001 | *phnA* | anthranilate synthase component I | 4.316 | 0.000 |
| PA1002 | *phnB* | anthranilate synthase component II | 4.184 | 0.000 |
| PA1126 | PA1126 | hypothetical protein | 1.561 | 0.000 |
| PA1127 | *gsp69* | probable oxidoreductase | 1.037 | 0.000 |
| PA1249 | *aprA* | alkaline metalloproteinase precursor | 1.007 | 0.000 |
| PA1871 | *lasA* | LasA protease precursor | 2.473 | 0.000 |

| | | | | |
|---|---|---|---|---|
| PA1900 | *phzB2* | probable phenazine biosynthesis protein | 3.285 | 0.000 |
| PA1901 | *phzC2* | phenazine biosynthesis protein PhzC | 1.223 | 0.000 |
| PA1903 | *phzZ2* | phenazine biosynthesis protein PhzE | 2.097 | 0.000 |
| PA1904 | *phzF2* | probable phenazine biosynthesis protein | 2.342 | 0.000 |
| PA2069 | PA2069 | probable carbamoyl transferase | 2.145 | 0.000 |
| PA2152 | PA2152 | probable trehalose synthase | 1.289 | 0.000 |
| PA2293 | PA2293 | hypothetical protein | 3.810 | 0.000 |
| PA2294 | PA2294 | probable ATP-binding component of ABC transporter | 6.190 | 0.000 |
| PA2295 | PA2295 | probable permease of ABC transporter | 16.416 | 0.000 |
| PA2296 | PA2296 | hypothetical protein | 4.514 | 0.000 |
| PA2297 | PA2297 | probable ferredoxin | 2.815 | 0.000 |
| PA2298 | PA2298 | probable oxidoreductase | 2.413 | 0.000 |
| PA2299 | PA2299 | probable transcriptional regulator | 2.031 | 0.000 |
| PA2300 | *chiC* | chitinase | 2.204 | 0.000 |
| PA2566 | PA2566 | conserved hypothetical protein | 1.925 | 0.000 |
| PA2588 | *vqsM* | probable transcriptional regulator | 1.079 | 0.000 |
| PA3137 | PA3137 | probable major facilitator superfamily (MFS) transporter | 3.148 | 0.001 |
| PA3361 | *lecB* | fucose-binding lectin PA-IIL | 1.097 | 0.000 |
| PA3477 | *rhlR* | transcriptional regulator RhlR | 0.704 | 0.000 |
| PA3478 | *rhlB* | rhamnosyltransferase chain B | 1.347 | 0.000 |
| PA3479 | *rhlA* | rhamnosyltransferase 1 subunit A | 1.065 | 0.000 |
| PA3724 | *lasB* | elastase LasB | 0.776 | 0.001 |

| | | | | |
|---|---|---|---|---|
| PA3808 | *yfhJ* | conserved hypothetical protein | 1.571 | 0.000 |
| PA4078 | PA4078 | probable nonribosomal peptide synthetase | 2.157 | 0.000 |
| PA4142 | PA4142 | probable secretion protein | 2.611 | 0.000 |
| PA4143 | *cyaB* | probable toxin transporter | 2.554 | 0.000 |
| PA4144 | *opmK* | probable outer membrane protein precursor | 3.677 | 0.000 |
| PA4214 | *phzE1* | phenazine biosynthesis protein PhzE | 2.097 | 0.000 |
| PA4215 | *phzF1* | probable phenazine biosynthesis protein | 2.342 | 0.000 |
| PA4216 | *phzG1* | probable pyridoxamine 5'-phosphate oxidase | 1.860 | 0.000 |
| PA4705 | *phuW* | PhuW | 1.349 | 0.001 |
| PA4834 | *cntl* | Cntl | 1.192 | 0.000 |
| PA5172 | *arcB* | ornithine carbamoyltransferase, catabolic | 1.030 | 0.000 |
| PA5173 | *arcC* | carbamate kinase | 1.450 | 0.000 |
| PA5534 | PA5534 | hypothetical protein | 1.177 | 0.000 |
| PA5535 | PA5535 | conserved hypothetical protein | 1.079 | 0.000 |
| PA5536 | *dksA2* | DksA2 | 1.655 | 0.000 |
| PA5538 | *amiA* | N-acetylmuramoyl-L-alanine amidase | 1.512 | 0.000 |
| PA5539 | PA5539 | hypothetical protein | 2.096 | 0.000 |
| PA5540 | PA5540 | hypothetical protein | 1.298 | 0.000 |
| **Down-regulated genes** | | | | |
| PA0041 | PA0041 | probable hemagglutinin | -1.244 | 0.000 |
| PA0314 | *fliY* | L-cysteine transporter of ABC system FliY | -1.965 | 0.001 |
| PA0905 | *rsmA* | translational regulator | -1.174 | 0.000 |
| PA1714 | *exsD* | ExsD | -0.955 | 0.041 |

| | | | | |
|---|---|---|---|---|
| PA1964 | *ybiT* | probable ATP-binding component of ABC transporter | -1.240 | 0.000 |
| PA2037 | PA2037 | hypothetical protein | -1.201 | 0.000 |
| PA2040 | *pauA4* | glutamylpolyamine synthetase | -1.498 | 0.000 |
| PA2309 | PA2309 | hypothetical protein | -2.039 | 0.001 |
| PA2310 | PA2310 | hypothetical protein | -1.496 | 0.000 |
| PA2311 | PA2311 | hypothetical protein | -16.716 | 0.000 |
| PA2675 | *hplT* | probable type II secretion system protein | -14.961 | 0.001 |
| PA2968 | *fabD* | malonyl-CoA-[acyl-carrier-protein] transacylase | -2.794 | 0.001 |
| PA3446 | *ssuE* | conserved hypothetical protein | -1.183 | 0.000 |
| PA3865.1 | PA3865.1 | pyocin S4 immunity protein | -1.120 | 0.000 |
| PA3895 | PA3895 | probable transcriptional regulator | -3.393 | 0.000 |
| PA4022 | *hdhA* | hydrazone dehydrogenase, HdhA | -2.190 | 0.000 |
| PA4155 | PA4155 | hypothetical protein | -1.261 | 0.000 |
| PA4292 | PA4292 | probable phosphate transporter | -1.421 | 0.000 |
| PA4315 | *mvaT* | transcriptional regulator MvaT, P16 subunit | -1.079 | 0.000 |
| PA4691 | PA4691 | hypothetical protein | -15.104 | 0.000 |
| PA5042 | *pilO* | type 4 fimbrial biogenesis protein PilO | -1.510 | 0.000 |
| PA5255 | *algQ* | alginate regulatory protein AlgQ | -0.395 | 0.041 |
| PA5264 | PA5264 | hypothetical protein | -1.393 | 0.000 |
| PA5383 | PA5383 | conserved hypothetical protein | -1.236 | 0.000 |
